# Malaria parasites coordinate host cell remodeling through the AP2-HCR DNA-binding protein

**DOI:** 10.64898/2026.06.23.733580

**Authors:** Amit Kumar Subudhi, Dhiviya Vedagiri, Sara Mfarrej, Reem Abu-Shamma, Rohit Satyam, Rahul P. Salunke, Di Liu, Maya Ayach, Ashraf Dada, Abdullah Fuaad Kadamany, David J.P. Ferguson, Sabrina Absalon, Arnab Pain

**Affiliations:** Pathogen Genomics Group, BioMedical Sciences (BioMed) Division, King Abdullah University of Science and Technology, Thuwal, Kingdom of Saudi Arabia; KAUST Core Labs, King Abdullah University of Science and Technology, Thuwal, Kingdom of Saudi Arabia; Department of Pathology and Laboratory Medicine, King Faisal Specialist Hospital Research Center, Jeddah, Kingdom of Saudi Arabia; College of Medicine, Al Faisal University, Riyadh, Saudi Arabia; Department of Biological and Medical Sciences, Faculty of Health and Life Sciences, Oxford Brookes University, Oxford, UK; Department of Biochemistry, Molecular Biology and Pharmacology, Indiana University School of Medicine, Indianapolis, Indiana, United States of America; International Institute for Zoonosis Control (IIZC), Institute for Vaccine Research and Development (IVReD)), Hokkaido University, Sapporo, Japan

## Abstract

Parasite-induced host cell remodeling is essential for *Plasmodium falciparum* survival during the intraerythrocytic developmental cycle (IDC), yet the transcriptional mechanisms coordinating this process remain poorly understood. Here, we characterize an essential ApiAP2 transcription factor, PfAP2-HCR (host cell remodeling), during both the asexual and sexual blood stages of *P. falciparum*. Conditional truncation of *pfap2-hcr* demonstrated that PfAP2-HCR is essential for ring stage development and parasite maturation. Loss of PfAP2-HCR disrupted the expression of host cell remodeling genes, resulting in the absence of Maurer’s clefts and collapse of the parasite protein export machinery. Integrated RNA-seq and ChIP-seq analyses revealed that PfAP2-HCR directly or indirectly regulates approximately 30% of exported proteins, including 27 essential exported factors, the parasitophorous vacuole membrane (PVM) resident protein EXP1, and the core PTEX component EXP2. We further show that PfAP2-HCR is also indispensable for gametocytogenesis, as conditional truncation results in either complete ablation or the formation of severely deformed gametocytes. Together, these findings establish PfAP2-HCR as a central transcriptional regulator that orchestrates the expression of host cell remodeling factors required to transform a terminally differentiated erythrocyte into a permissive environment for parasite survival, development, and transmission.

## Main

All clinical symptoms of malaria arise during the asexual blood stage development. Merozoites invade red blood cells (RBCs) and develop within a parasitophorous vacuole (PV) through ring, trophozoite, and schizont stages before segmentation releases 16–32 daughter merozoites that invade new RBCs^1^. Throughout this cycle, parasites remodel the infected RBC (iRBC) to ensure nutrient uptake, cytoadherence, and immune evasion^2^. Host-cell remodeling is not restricted to asexual blood stages; it also occurs during gametocyte development^3^ and during liver-stage development following invasion of sporozoites, parasites reorganize hepatocytes to support intracellular growth^4,5^. Thus, remodeling of diverse host-cell environments represents a central adaptive strategy used by malaria parasites to survive, replicate and transmit.

During asexual blood stage development, following invasion, the parasite rapidly establishes the parasitophorous vacuole membrane (PVM) and initiates extensive host cell remodeling by exporting hundreds of effector proteins across it. This transport is mediated by the parasite-derived protein conducting channel known as the *Plasmodium* translocon of exported proteins (PTEX), whose three core components are EXP2, PTEX150, and HSP101^6–8^. Conditional disruption of any core PTEX component or chemical inhibition of protein export results in parasite death, confirming its essentiality^9–11^.

Parasite exported proteins (exportome) can be divided into two groups based on whether they contain a *Plasmodium* export element (PEXEL)^12^, also known as the vacuolar translocation sequence (VTS)^13^. Exported proteins that do not contain PEXEL motifs are called PEXEL-negative export elements (PNEPs)^14^. The *P. falciparum* exportome includes ∼460 PEXEL-positive and >40 PNEPs, of which at least 71 are putatively essential for survival^15^. The exportome can further be divided into two classes of proteins: (1) proteins that are expressed during the early ring stage and are necessary for establishing the tubovesicular network (i.e. skeleton binding protein 1, ring exported protein 1-2, membrane-associated histidine-rich protein 1, *P. falciparum* erythrocyte membrane protein 3) and (2) proteins that are expressed later during the late ring or early trophozoite stage and perform other functions, such as antigenic variation and cell signalling (i.e. FIKK kinases, PfEMP1, RIFINS and STEVORS)^2,16,17^. The expression of the majority of these genes is initiated in the late schizont stage, continues throughout the invasion process and peaks during the early ring stage ^17^. These genes need to be coexpressed in a short duration of time during the late-schizont, merozoite and early ring stages, which must be under the control of a common gene-regulatory complex.

*P. falciparum* exhibits tightly regulated, stage-specific transcription, with roughly half of all genes expressed immediately before their proteins are required^17,18^. Gene expression is modulated by a combination of sequence-specific transcription factors, epigenetic control, and chromatin organization^19,20^. The parasite genome encodes few canonical transcription factors; instead, regulation relies largely on the ∼30 Apicomplexan Apetala2 (ApiAP2) family of DNA-binding proteins present across apicomplexan parasites^21,22^.

Individual ApiAP2 factors orchestrate key developmental transitions and stress responses: AP2-G and its derivatives govern sexual commitment and gametocytogenesis^23–25^, AP2-O and AP2-Z proteins control ookinete maturation^26–28^, AP2-I regulates invasion genes^29^, AP2-HS mediates heat-shock responses^30^, and AP2-P regulates multiple pathogenic processes^31^. The combinatorial action of these transcription factors provides the temporal precision necessary for rapid adaptation within the host.

Despite advances in defining ApiAP2 functions, the transcriptional program governing early ring stage remodeling remains largely unknown. Because many exported proteins and PTEX components are expressed during invasion, we hypothesized that a dedicated ApiAP2 factor coordinates this gene network.

Here, we identify PfAP2-HCR (*<u>h</u>ost <u>c</u>ell <u>r</u>emodeling*), an ApiAP2 DNA-binding protein essential during ring stage development. Using conditional gene truncation, high-resolution phenotyping via ultrastructure expansion microscopy (U-ExM), transcriptomics, chromatin occupancy, and proteomics analyses, we demonstrate that PfAP2-HCR directly regulates a broad subset of exported proteins that are crucial for remodeling infected RBCs and parasite survival. We also show that PfAP2-HCR is also indispensable for gametocytogenesis, extending its function beyond asexual blood-stage development. This work establishes PfAP2-HCR as a central transcriptional regulator linking gene-expression control to functional host-cell modification, thereby defining a previously unrecognized gene-expression network underlying malaria parasite development within the red cells and pathogenesis.

## Results

### PfAP2-HCR is essential for ring stage development and gametocytogenesis

In a recent study^31^, we characterized the ApiAP2 transcription factor PfAP2-P, a DNA-binding protein that modulates multiple pathogenic processes through its circadian-like, bimodal expression pattern during the IDC. PfAP2-P exhibits two major expression peaks, at approximately 16- and 40-hours post-invasion (h.p.i.), and directly regulates nine *apiap2* genes across these phases, three of which remain uncharacterized (*PF3D7_0420300*, *PF3D7_0613800* and *PF3D7_1115500*) (**Extended Data Fig. 1a**). Conditional truncation of *pfap2-p* resulted in the downregulation of two and upregulation of one *apiap2* genes around 40 h.p.i. (**Extended Data Fig. 1a**).

Of the two downregulated *apiap2s*, *PF3D7_0420300*, hereafter designated PfAP2-HCR, showed a pronounced expression peak immediately following that of *pfap2-p* during the IDC, consistent with direct transcriptional activation by PfAP2-P (**Extended Data Fig. 1b**). As PfAP2-HCR is highly expressed during the late schizont stage and shows elevated transcript levels in merozoite and ring stages of the intraerythrocytic developmental cycle^31,32^, as well as in ookinete and sporozoite stages^33–36^, we hypothesized that it contributes to transcriptional control during these stages (**Extended Data Fig. 2a**). Supporting this notion, proteomic and co-immunoprecipitation studies from our group and others have identified PfAP2-HCR in association with chromatin-modifying and transcriptional regulators including PfAP2-P^31^, PfMORC^37,38^, PfISWI^39^, and PfSET10 ^40^, suggesting that it participates in multi-protein gene-regulatory complexes.

*PfAP2-HCR* is a large, single-exon gene (10,422 bp excluding 5′ and 3′ UTRs) encoding a 3,473–amino acid protein that contains two AP2 DNA-binding domains located at residues 1,959–2,008 and 3,266–3,317, respectively (**Fig. 1a**). Orthologues of PfAP2-HCR are conserved across most apicomplexan lineages but are absent in *Cryptosporidium* species and their more distantly related alveolate relatives *Chromera velia* and *Vitrella brassicaformis* (**Extended Data Fig. 2b**). Functional data indicate that PfAP2-HCR and its orthologues in *P. berghei* and *P. knowlesi* are essential for asexual blood-stage development ^41–45^ (**Extended Data Fig. 2c**).

**Fig. 1.**
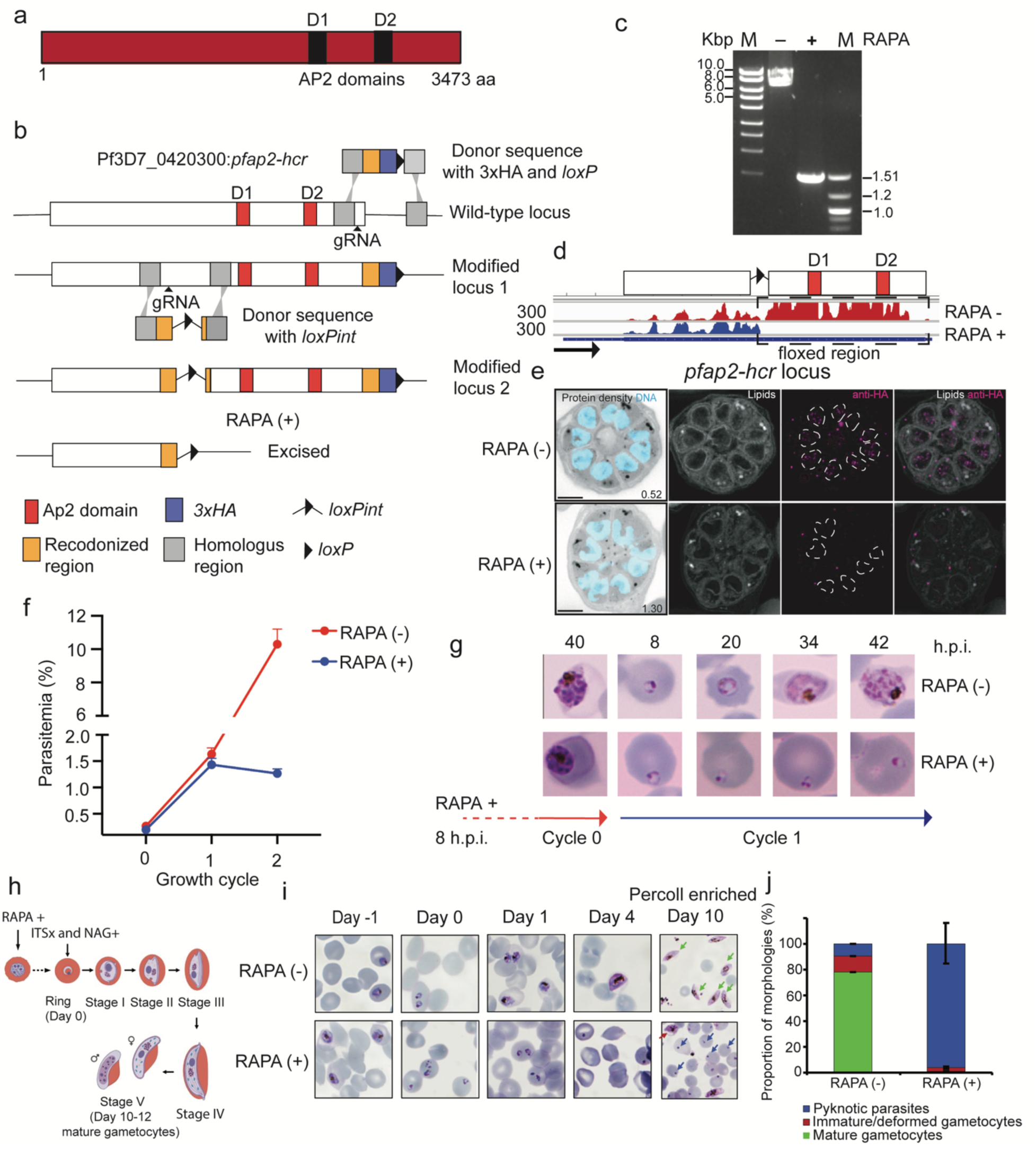
PfAP2-HCR is an asexual ring stage and gametocytogenesis essential AP2 DNA-binding protein. **a,** Domain organization of PfAP2-HCR showing the AP2 DNA-binding domains. **b**, Schematic representation of the conditional truncation strategy involving the excision of the *loxP*-flanked sequence encoding the AP2 DNA-binding domains. **c**, PCR-based confirmation of *loxP*-flanked sequence excision in rapamycin (RAPA)-treated parasites. **d**, RNA-seq read coverage plot demonstrating depletion of RNA reads corresponding to the excised region in RAPA-treated parasites. The arrow represents direction of transcription, and the numbers on the side represent read coverage. **e**, RAPA-treated (RAPA+) and control (RAPA−) parasites expressing PfAP2-HCR-3HA were prepared by U-ExM, stained with N-hydroxysuccinimide (NHS) ester (grayscale), BODIPY FL (white), SYTOX (cyan), and anti-HA antibodies (magenta) and imaged by Airyscan 2 microscopy. The white dashed line marks the nuclear envelope. Scale bars = 5 µm. **f**, Replication of control (RAPA−) and treated (RAPA+) parasites over three growth cycles (average parasitemia ± s.d.; *n*= 3 biological replicates). **g**, Images of Giemsa-stained parasites from cycle 0 through the beginning of cycle 1 following treatment with DMSO (RAPA–) or rapamycin (RAPA+) at ∼8 hours post-invasion (h.p.i.) in cycle 0. **h**, Schematic representation of the gametocytogenesis induction along with *pfap2-hcr* conditional truncation. **i**, Giemsa-stained microscopic images from day -1, 0, 1, 4 and percoll-enriched day 10 cultures from either control (RAPA−) and treated (RAPA+) parasites. **j**, Quantification of gametocytes (mature/immature/deformed) and pyknotic parasites from control (RAPA−) and treated (RAPA+) day 10 percoll-enriched parasites. Mature, deformed gametocytes and pyknotic parasites are highlighted in green, maroon and blue arrows, respectively. Error bars are representation of standard deviation from two independent experiments.

To investigate the function of PfAP2-HCR, we generated a triple human influenza hemagglutinin (3HA)-tagged PfAP2-HCR inducible AP2-domains knockout of *P. falciparum* (PfAP2-HCR-3HA:*loxP*) in a parental line expressing rapamycin (RAPA)-inducible dimerizable Cre recombinase (DiCre)^46^ (**Fig. 1b and Methods**). Upon RAPA treatment of tightly synchronized parasites at ∼5 h.p.i., efficient excision of the loxP-flanked AP2-domains encoding region was achieved by 40 h.p.i. within the same cycle, as confirmed by PCR and RNA-seq (**Fig. 1c–d**).

Through U-ExM, we also showed that mock-treated controls (DMSO-treated; RAPA −), PfAP2-HCR-3HA is readily detected within the parasite nucleus, whereas RAPA treatment (RAPA +) results in efficient loss of the nuclear PfAP2-HCR signal, consistent with loss of protein expression from the excised region (**Fig. 1e**). We next examined the expression of PfAP2-HCR across the intraerythrocytic developmental cycle using U-ExM. PfAP2-HCR protein was not detectable from the late ring stage until the onset of cytokinesis, consistent with transcriptomic data showing that *pfap2-hcr* RNA levels peak during schizont-stage development (**Extended Data Fig. 3a**).

To determine its essentiality during the IDC, we performed growth assay by adding RAPA/DMSO at ∼5 h.p.i. in cycle 0 and determined the growth of both mock-treated control (i.e. DMSO-treated) and RAPA-treated parasites for the next two growth cycles (cycle 0-2).

Conditional truncation of *pfap2-hcr* (*Δpfap2-hcr*) at cycle 0 showed that both control and *Δpfap2-hcr* parasites were able to proceed to the next cycle (i.e. cycle 1) without any significant difference in parasitemia between control and *Δpfap2-hcr* parasites suggesting that there were no issues with parasite egress and/or invasion (**Fig. 1f**). Interestingly, in contrast, control parasites proceeded to cycle 2 as expected whereas *pfap2-hcr* parasites failed to complete cycle 1 with a dramatic reduction in parasitemia compared with controls in cycle 2 (**Fig. 1f).**

To determine the developmental stage at which *Δpfap2-hcr* parasites were affected in cycle 1, we collected samples for light microscopy at multiple time points from the end of cycle 0 through the end of cycle 1 (**Fig. 1g**). Strikingly, *Δpfap2-hcr* parasites invaded successfully and formed early ring stage parasites in cycle 1 but failed to progress further and subsequently degenerated (**Fig. 1g**), indicating that PfAP2-HCR is essential for ring stage maturation and subsequent intraerythrocytic development.

To pinpoint the temporal requirement of PfAP2-HCR, RAPA was administered to synchronized cultures at distinct IDC stages—8, 24, and 36 h.p.i., corresponding to the ring, trophozoite, and early schizont stages, respectively (**Extended Data Fig. 3b**). As complete excision occurs within ∼24 h post-RAPA treatment^31^, this approach allowed stage-specific functional interrogation. In line with the initial experiment, truncation induced at 8 h.p.i. or 24 h.p.i. produced the same phenotype: parasites failed to progress beyond the ring stage in the next cycle (**Extended Data Fig. 3b**). In contrast, RAPA addition at 36 h.p.i. allowed normal development through the ensuing cycle 1 but resulted in growth arrest at the ring stage in cycle 2 (**Extended Data Fig. 3b**). Together, these results demonstrate that PfAP2-HCR is essential specifically during the ring stage of *P. falciparum* development.

Because parasite-induced host-cell remodeling also occurs during gametocytogenesis, we next tested whether PfAP2-HCR is required during sexual blood-stage development. We performed two complementary conditional-excision experiments (**Fig. 1h and Extended Data 4a**). In the first approach, rapamycin was added in the preceding asexual cycle, at the late trophozoite/early schizont stage (∼30 h.p.i.; day −1 relative to gametocyte induction), to induce earlier truncation of *pfap2-hcr* before sexual-stage development (**Fig. 1h**). In the second approach, rapamycin or DMSO was added on day 0 of gametocyte induction, at the ring/early committed gametocyte stage, together with N-acetyl glucosamine to eliminate asexual parasites and ITS-x to support gametocyte development (**Extended Data Fig. 4a**). Here, gametocyte induction is referred to as a process to eliminate asexual parasites by adding N-acetyl glucosamine and ITS-x to support growth of sexually committed parasites (Methods).

In both experimental conditions, rapamycin-treated parasites showed a pronounced defect in gametocyte formation and maturation (**Fig. 1i, j and Extended Data Fig. 4b, c).** By day 10 of gametocyte induction, *Δpfap2-hcr* cultures from the first condition contained predominantly pyknotic infected red blood cells, with only a small proportion of deformed gametocytes and almost no morphologically mature gametocytes (**Fig. 1j**). In the second condition, Δ*pfap2-hcr* cultures contained comparable proportions of pyknotic infected red blood cells and deformed gametocytes, together with only a minor fraction of mature gametocytes (**Extended Data Fig. 4c**). In contrast, DMSO-treated controls from both conditions progressed efficiently to morphologically mature stage IV/V gametocytes (**Fig. 1j and Extended Data Fig. 4c**). These results indicate that PfAP2-HCR is required not only during asexual blood-stage development but also for normal sexual blood-stage maturation. The phenotype was more severe when *pfap2-hcr* was excised in the preceding late trophozoite or early schizont stage, consistent with possible earlier disruption of the export machinery and expression of exported proteins during early gametocyte development (**Fig. 1j)**. A similar developmental arrest has been reported following conditional depletion of PfHSP101 during stage I gametocyte development^9^. By contrast, when excision was induced on day 0, some early gametocyte development may have proceeded before complete loss of PfAP2-HCR function, leading to formation of deformed immature gametocytes, suggesting a continued requirement for PfAP2-HCR-dependent host-cell remodeling factors during gametocytogenesis.

### PfAP2-HCR regulates the expression of exported proteins required for host cell remodeling

To investigate the transcriptional role of PfAP2-HCR, we performed bulk RNA-seq analyses on synchronized parasites following conditional truncation of *pfap2-hcr*. DMSO or RAPA was added at ∼5 h.p.i., and samples were collected at ∼45 h.p.i. in the same cycle and at ∼5 h.p.i. in the subsequent cycle (cycle 1) (**Fig. 2a**). These timepoints correspond to the peak expression of *pfap2-hcr* during the IDC, providing maximal sensitivity for the detection of its regulatory targets.

**Fig. 2.**
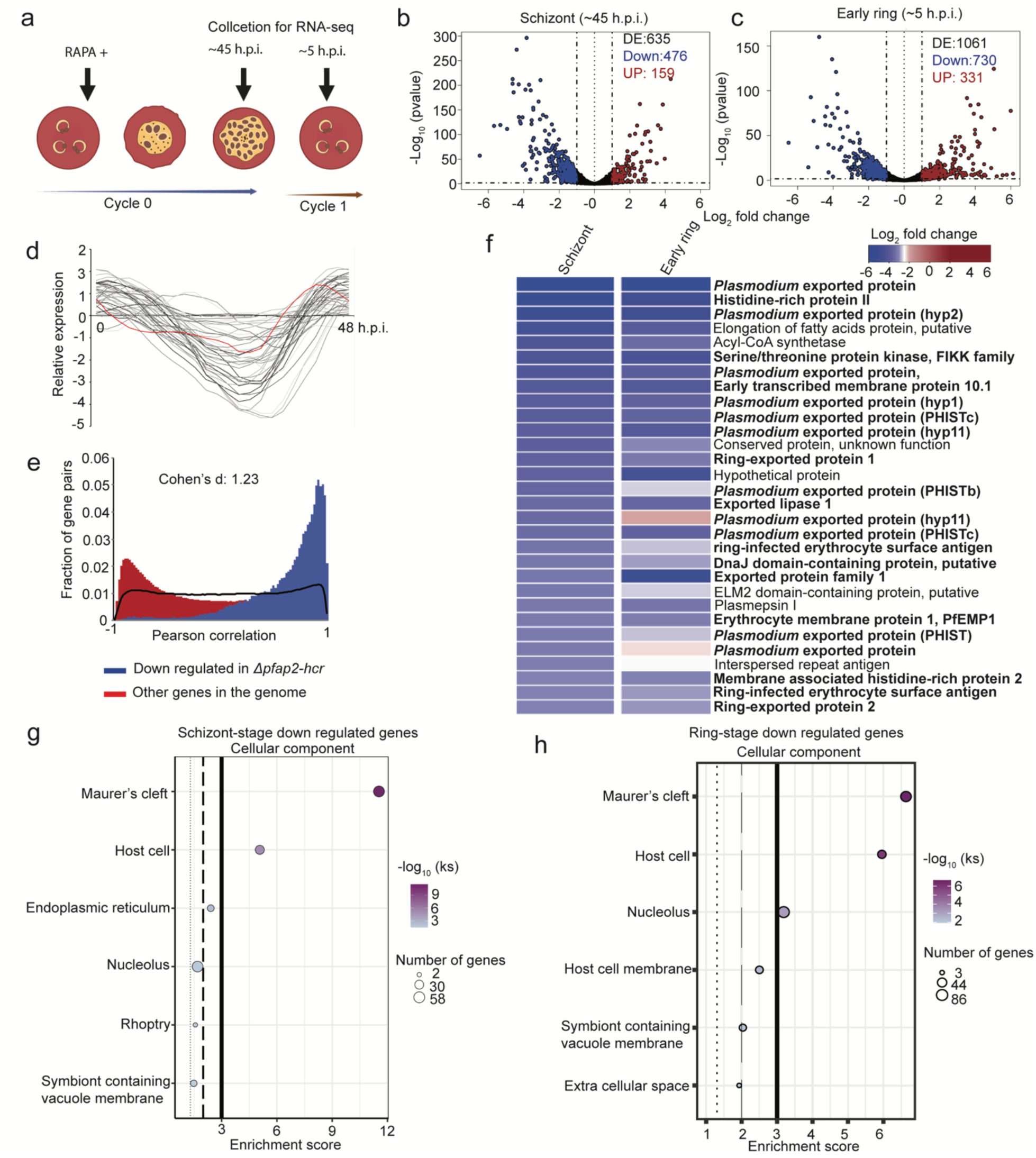
PfAP2-HCR regulates a broad set of exported proteins required for host cell remodeling. **a**, Schematic representation of the sampling time points for control (RAPA–) and rapamycin-treated (RAPA+) parasites collected for RNA-seq analysis. **b,c**, Volcano plots showing differentially expressed genes in *Δpfap2-hcr* parasites at ∼45 h post-invasion (h.p.i.) in cycle 0 (**b**) and ∼5 h.p.i. in cycle 1 (**c**). Upregulated genes (log₂(FC) > 1, *P* < 0.05) are shown in maroon, and downregulated genes (log₂(FC) < -1, *P* < 0.05) are shown in blue. **d**, Expression pattern of the 50 most significantly downregulated genes in *Δpfap2-hcr* schizont-stage parasites and *pfap2- hcr* (red line) during the intraerythrocytic developmental cycle^16^. **e**, The distribution of coexpression values, as measured by the Pearson correlation coefficients, are plotted for two gene pair categories; coexpression correlation between down-regulated genes in *Δpfap2-hcr* parasites (blue) and coexpression correlation between all other genes (maroon). **f**, The 30 most significantly downregulated genes in *Δpfap2-hcr* schizont-stage parasites. Genes encoding exported proteins are highlighted in bold text. **g-h**, Bubble plots showing enriched Gene Ontology cellular component (CC) terms among significantly downregulated genes (log₂(FC) < - 1, *Padj* < 0.05) in schizonts (**g**) and early ring stage (**h**) of the *Δpfap2-hcr* parasites.

Comparative transcriptomic analysis between Δ*pfap2-hcr* and control parasites revealed 635 and 1,061 significantly differentially expressed genes (adjusted *P* < 0.05) at ∼45 h.p.i. and ∼5 h.p.i., respectively (**Fig. 2b,c; Supplementary Data 1**). In both datasets, the majority of affected genes were downregulated upon PfAP2-HCR truncation, suggesting PfAP2-HCR functions primarily as a transcriptional activator (**Fig. 2b,c**).

Expression profiling of the 50 most strongly downregulated genes in Δ*pfap2-hcr* parasites showed that their transcript levels peak immediately after *pfap2-hcr* expression during the IDC^16^ (**Fig. 2d**), suggesting direct transcriptional control by PfAP2-HCR. Supporting this, a pairwise correlation analysis revealed that these downregulated genes exhibit markedly higher co-expression than the genome-wide background (Cohen’s *d* = 1.23), indicative of coordinated regulation by a common transcriptional regulator—PfAP2-HCR (**Fig. 2e**).

Among the 30 most strongly downregulated genes, 23 encode exported proteins predicted to mediate parasite-induced remodeling of the infected red blood cells (iRBCs) (**Fig. 2f**; **Supplementary Data 1**). Gene Ontology (GO) enrichment analysis of downregulated genes in schizont- and ring stage Δ*pfap2-hcr* parasites revealed strong enrichment for the cellular component terms Maurer’s cleft, host cell, and host cell membrane (**Fig. 2g,h**). These findings are consistent with the observed developmental arrest of Δ*pfap2-hcr* parasites at the ring stage, suggesting that PfAP2-HCR controls the expression of genes essential for host cell remodeling and subsequent intraerythrocytic development.

To assess the global impact of PfAP2-HCR on the exported protein repertoire, we analyzed transcriptomic changes across the exportome—comprising all proteins predicted or experimentally validated to be exported^15^. Of the 305 exported proteins (excluding *var*, *rifin*, and *stevor* families), 71 (23%) were significantly downregulated upon *pfap2-hcr* truncation, indicating that the loss of AP2 domains in PfAP2-HCR perturbs a substantial fraction of the exportome (**Supplementary Data 1**). The affected genes include several key determinants of host cell remodeling and trafficking, such as *resa* and *resa3*, *hsp40*, *hrp2*, *rex1–3*, members of the *epf1/3/4* family, *FIKK9.4*, *FIKK9.6*, *FIKK12*, *ptp1–6*, *etramp10.2* and *etramp14*, and *mahrp1/2*. Additionally, transcripts encoding the parasitophorous vacuole membrane (PVM) proteins *exp1* and *exp2*—the latter of which is a core component of the PTEX translocon—were significantly reduced, whereas other PTEX constituents (*hsp101* and *ptex150*) remained unaffected. GO enrichment of the full downregulated set further identified perturbations in the signal transduction, lipid metabolism, and heme catabolism pathways (**Extended Data Fig. 5a,b**). Together, these results demonstrate that PfAP2-HCR is a major transcriptional regulator of host cell remodeling and export-related processes.

To examine the stage-specific expression patterns of the affected exportome, we performed gene expression clustering of all known exported proteins across the IDC, identifying four distinct temporal clusters (**Extended Data Fig. 6a and Supplementary Data 2**). Cluster 1 comprises genes expressed at early schizont stages, Cluster 2 those with near-constant expression across the IDC, Cluster 3 genes peaking at late schizont and early ring stages, and Cluster 4 genes upregulated during trophozoite development. Downregulated genes in Δ*pfap2-hcr* were highly enriched in Cluster 3 compared to all other clusters (two-sided Fisher’s exact test, odds ratio = 17.34, *p* = 2.85 × 10⁻¹⁶; **Extended Data Fig. 6b**). Cluster 3 genes, such as *resa*, *rexs*, *etramps*, *sbp1*, a few *fikks*, and *ptp* family members, encode exported proteins that prepare the parasite for remodeling of the host cell immediately after invasion of the red cells, underscoring PfAP2-HCR’s role in regulating this critical developmental transition.

GO enrichment of upregulated genes identified adhesion of symbiont to host cell (antigenic variation) as the most significantly enriched biological process (**Extended Data Fig. 5c,d**). Genes contributing to this term included several *rifin* and *stevor* family members (**Supplementary Data 1**).

### PfAP2-HCR directly regulates genes required for host cell remodeling

To identify the direct genomic targets of PfAP2-HCR, we performed chromatin immunoprecipitation followed by sequencing (ChIP–seq) using parasites expressing 3×HA-tagged PfAP2-HCR. Because PfAP2-HCR is highly expressed during the late schizont and merozoite stages, ChIP–seq was performed on compound 2-treated (an inhibitor of egress^47^) *P. falciparum* culture containing very late segmented schizonts to capture the full spectrum of its DNA-binding activity (**Fig. 3a**). Across two biological replicates, we identified 542 peaks with shared genomic occupancy (Methods), of which 303 (55%) were located in intergenic or promoter regions upstream of at least one gene—most within 2 kb of the translational start site (**Fig. 3b,c and Supplementary Data 3**).

**Fig. 3.**
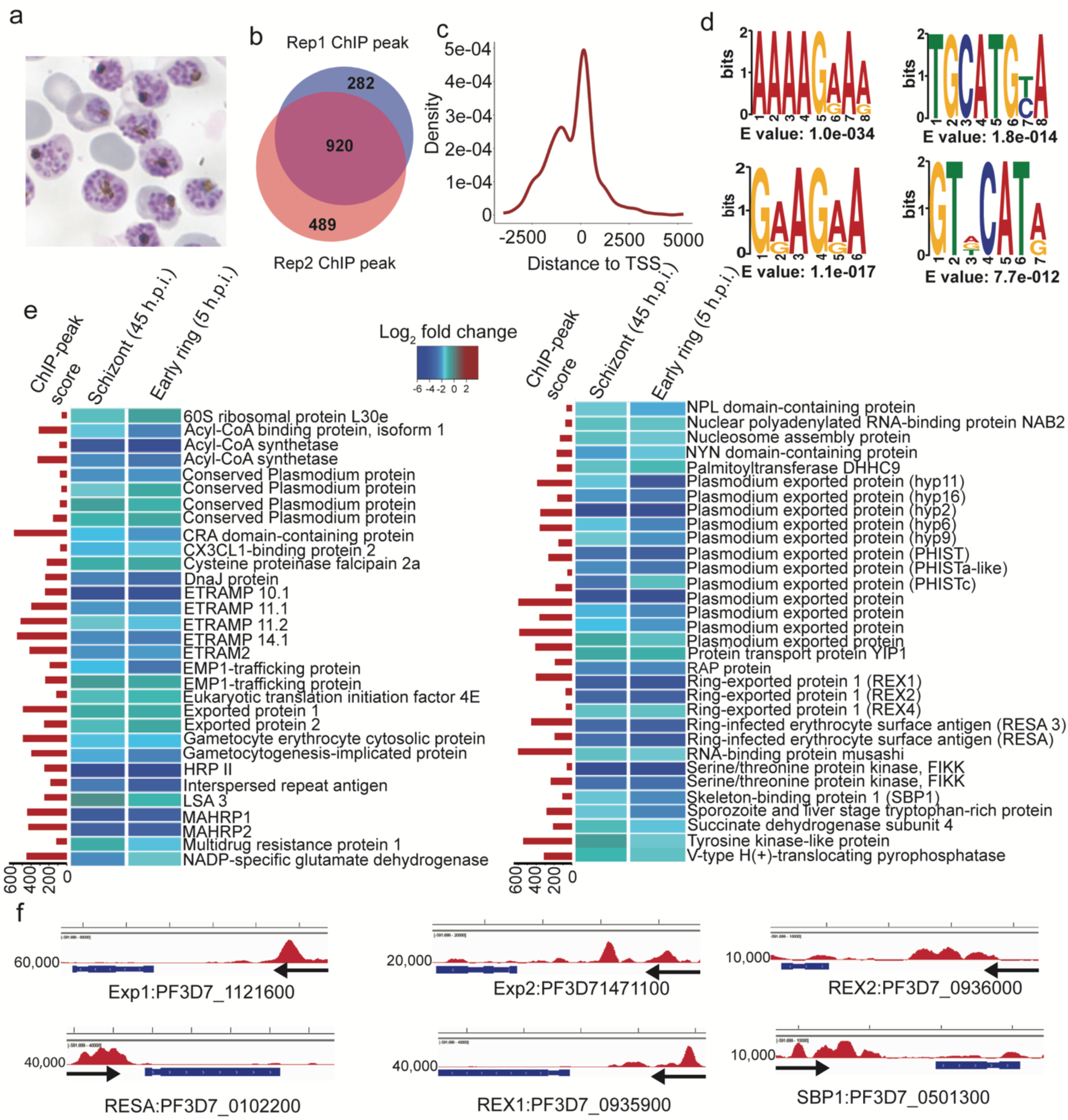
PfAP2-HCR directly regulates exported proteins required for host cell remodeling. **a**, Image of Giemsa stained C2-treated parasites used for ChIP–seq experiments to identify the genomic occupancy of PfAP2-HCR. **b**, Venn diagram comparing PfAP2-HCR promoter peaks identified from two biological replicates using *bedtools*. **c**, Distribution of ChIP–seq peak summits relative to predicted translational start sites (TSSs). **d**, The four most significantly enriched DNA motifs bound by PfAP2-HCR. **e**, Representative genes from all the genes directly and positively regulated by PfAP2-HCR. Bar plots (left) show ChIP peak scores from two biological replicates determined using *bedtools intersect*, whereas heatmaps (right) display the expression levels (log₂ fold change) of the corresponding genes. **f**, PfAP2-HCR occupancy at the putative promoter regions of representative iRBC remodeling genes. Input-subtracted ChIP track is shown, with enrichment scores (left) and transcriptional direction (arrows) indicated.

Motif enrichment analysis of PfAP2-HCR–bound regions revealed several significantly enriched DNA motifs, including AAAAGRAR (*E* = 1.0 × 10⁻³⁴), GRAGRA (*E* = 2.2 × 10⁻¹⁹), TGCATGYA (*E* = 9.5 × 10⁻¹⁵), and GTDCATR (*E* = 6.3 × 10⁻¹⁷) (**Fig. 3d**). The GTDCATR motif closely resembles the binding motifs of PfAP2-I and PfAP2-P (*P* = 4.61 × 10⁻³), consistent with previous evidence that PfAP2-HCR associates with these transcription factors and the chromatin remodeling proteins PfMORC and PfISWI as part of a multiprotein regulatory complex ^29,31,38,39,48^. Previous work using protein-binding microarrays and purified N-terminal glutathione-S-transferase (GST)–tagged DNA-binding domains of multiple PfAP2 proteins, including PfAP2-HCR, reported that the first AP2 DNA-binding domain of PfAP2-HCR preferentially binds a CACACA motif^49^. Notably, this motif differs from the binding motifs identified in this study. This discrepancy may reflect differences between *in vitro* and *in vivo* binding contexts. The earlier study examined an isolated DNA-binding domain under in vitro conditions, whereas our analysis captures genome-wide binding of the full-length protein in the parasite nucleus. *In vivo*, PfAP2-HCR is likely to function within multi-protein regulatory complexes and to bind DNA within a chromatin environment whose accessibility and epigenetic features may further influence binding specificity.

PfAP2-HCR binding was highly enriched at the promoters of genes encoding exported proteins, corroborating the transcriptional downregulation observed in Δ*pfap2-hcr* parasites. Gene ontology analysis of bound targets revealed significant enrichment for cellular components associated with the infected erythrocyte surface, Maurer’s clefts, heterochromatin, and the PTEX translocon (**Extended Data Fig. 7**). PfAP2-HCR bound to the promoters of several exported protein genes that were transcriptionally downregulated upon *pfap2-hcr* truncation, including *rex1/2/4*, *resa* and *resa3*, *fikk* family kinases, *etramp* genes, *emp1*-trafficking proteins, *mahrp1/2*, *exp1/2*, and *hrp2* (**Fig. 3e,f and Supplementary Data 3**). In total, 51 exported protein–encoding genes (excluding antigenic variation families) were directly bound by PfAP2-HCR, supporting its role as a major transcriptional regulator of parasite proteins involved in host cell remodeling.

PfAP2-HCR also bound to the promoter regions of 16 *var* genes, which encode variant surface antigens and are typically positioned in subtelomeric heterochromatic regions. However, no binding was detected at promoters of other antigenic variation gene families such as *rifins*, *stevors*, or *pfmc-2tms*, indicating a preference for subtelomeric *var* promoters. Conditional deletion of PfAP2-HCR led to the downregulation of at least 15 *var* genes during the ring and late schizont stages (**Supplementary Data 1**), although none of these genes were directly bound by PfAP2-HCR. As *vars* do not express during the stages when sampling was performed for RNA-seq and ChIP-seq, we believe that the observed downregulation of *vars* was likely an indirect or bystander effect of *pfap2-hcr* deletion rather than a consequence of direct transcriptional control.

Interestingly, PfAP2-HCR also bound the putative promoters of 10 other *apiap2* genes, including *ap2-l*, *ap2-exp*, *ap2-g*, *ap2-hs, ap2-o*, *ap2-o3*, and *ap2-o5*, as well as other uncharacterized members (*PF3D7_0934400* and *PF3D7_1115500*), and its own promoter, suggesting autoregulation (**Extended Data Fig. 8 and Supplementary Data 3**). Consistent with previous reports that multiple ApiAP2 transcription factors form combinatorial regulatory complexes where they regulate each other ^29,31^, these findings indicate that PfAP2-HCR functions within an integrated ApiAP2 network that coordinates gene expression across key developmental stages of the parasite. Interestingly, conditional truncation of *pfap2-hcr* resulted in significant upregulation of nearly all *apiap2* genes whose promoters were bound by PfAP2-HCR, including *pfap2-i*, *pfap2-g*, *pfap2-exp*, *pfap2-hs*, *pfap2-p* and *pfap2-fg* (adjusted *P* < 0.05; log₂ fold-change > 1) (**Supplementary Data 1)**. This effect was observed at both late schizont and early ring stages, suggesting that PfAP2-HCR as part of the repressive regulatory complex^31,38,40,48^ that binds to the promoter of *ap2s* represses a subset of ApiAP2 transcription factors during these developmental windows.

To explore the genome-wide occupancy of PfAP2-HCR in other blood-stages, we performed ChIP-seq at 20, 30 and 40 h.p.i. As the expression of *pfap2-hcr* is limited to late schizont, merozoite and early ring stages and its essentiality during late ring or early trophozoite stage, we anticipate its limited regulatory role during other IDC stages. Complementing this observation, we only observed 30, 38 and 2 ChIP-seq peaks at 20, 30 and 40 h.p.i respectively, suggesting there is no or limited role during other IDC stages (**Supplementary Data 4)**.

### PfAP2-HCR is a transcriptional regulator of essential exported proteins required for host cell remodeling

Integration of RNA-seq and ChIP–seq data identified 93 of 305 (30%) known or predicted exported proteins (both PEXEL-positive and PNEPs) as directly or indirectly regulated by PfAP2-HCR, indicating its role as a key regulator of genes essential for host cell remodeling and parasite survival (**Fig. 4**). To assess which of these targets are indispensable, we compiled a list of exported genes reported to be essential for parasite viability from targeted gene disruption studies and large-scale piggyBac transposon mutagenesis screening ^15^. Among the 71 essential exported proteins (PEXEL and PNEPs) curated from these datasets, 27 (38%; 12 directly and 15 indirectly) were regulated by PfAP2-HCR, indicating that this transcription factor controls a substantial proportion of the essential exportome during the IDC.

**Fig. 4.**
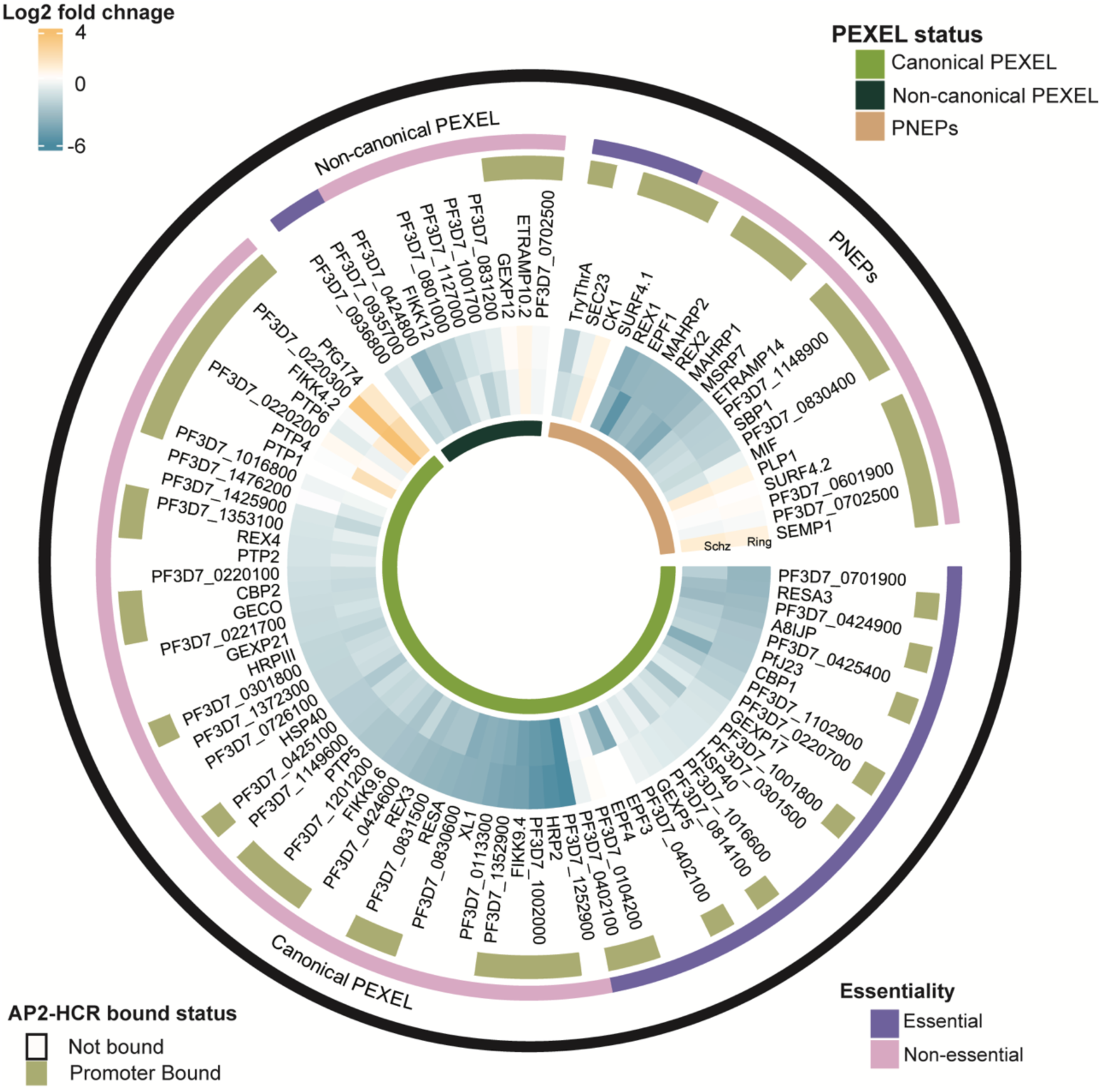
PfAP2-HCR regulates the transcriptional network controlling host cell remodeling during the early stages of parasite development within the RBCs. Circos plot showing the expression status of PfAP2-HCR-regulated exported proteins (both essential and non-essential) in RAPA-treated (*Δpfap2-hcr*) parasites compared to controls. Genes directly or indirectly regulated by PfAP2-HCR were identified based on PfAP2-HCR binding to their promoter regions. PNEPs, PEXEL-negative exported proteins. The refined list of exported proteins and their essentiality status was obtained from Jónsdóttir *et al.* (2021).^15^.

### PfAP2-HCR associates with other regulatory proteins

To identify proteins that interact with PfAP2-HCR, we performed chromatin immunoprecipitation followed by mass spectrometry (ChIP–MS) using PfAP2-HCR–3HA parasites. This analysis revealed several known and putative transcription factors and chromatin remodelers, including PfAP2-P, PfAP2-EXP, PfAP2-O, the general transcription factor TF-BTF3, the microrchidia family protein PfMORC, and a chromodomain-helicase-DNA-binding protein 1 (CHD1) homolog (**Extended Data Fig. 9; Supplementary Data 5**). These findings indicate that, similar to other ApiAP2 family members, PfAP2-HCR functions as part of a multi-protein regulatory complex comprising both transcriptional regulators and chromatin-remodeling factors. The detection of PfAP2-P and PfMORC within the complex is consistent with previous reports of their physical and functional association with PfAP2-HCR^31,38^.

### Infected erythrocytes harboring *Δpfap2-hcr* parasites display hallmarks of defective host cell remodeling

To examine the functional consequences of PfAP2-HCR loss on host cell remodeling, we analyzed Δ*pfap2-hcr* parasites by transmission electron microscopy at 12 and 24 h post-invasion in cycle 1 (following RAPA treatment at early ring stage in cycle 0). To identify any developmental difference a random sample of between 50 and 80 parasites were recorded for each sample and any difference was quantified. The developmental stages parasite development were characterised as firstly rings which are elongated or crescentic-shaped (**Fig. 5a**), then early trophozoites which were more spherical shape with a central nucleus (**Fig. 5b** and **c**) and then late trophozoites containing a food vacuole with heamozoin crystals (**Fig. 5d**). In addition, in the mock-treated control groups a number infected RBC at 24 hrs exhibited flattened vacuoles running parallel to the RBC plasma membrane - Mauer’s clefts (**Fig. 5d insert**). In both mock-treated controls and RAPA+ samples at 12 h.p.i, the majority of parasites were at the ring or early trophozoite stage (90+%). However, divergence between the groups was noted at 24 h.p.i. While there was little evidence of parasite development in the RAPA+ samples between 12 and 24 h.p.i. with the proportion of rings or early trophozoites remaining similarly high (90% vs 89%). In contrast, in the mock-treated control samples at 24 h.p.i a significant proportion (38%) had advanced to the later stage with food vacuoles containing hemozoin crystals. While Maurer’s clefts, hallmarks of host cell remodeling, were clearly visible in infected erythrocytes of mock-treated controls, they were not observed in any TEM images of Δ*pfap2-hcr*–infected cells (**Fig. 5d inset**). Collectively, these molecular and ultrastructural data establish PfAP2-HCR as a central transcriptional regulator of exported proteins required for host cell remodeling and parasite development.

**Fig. 5.**
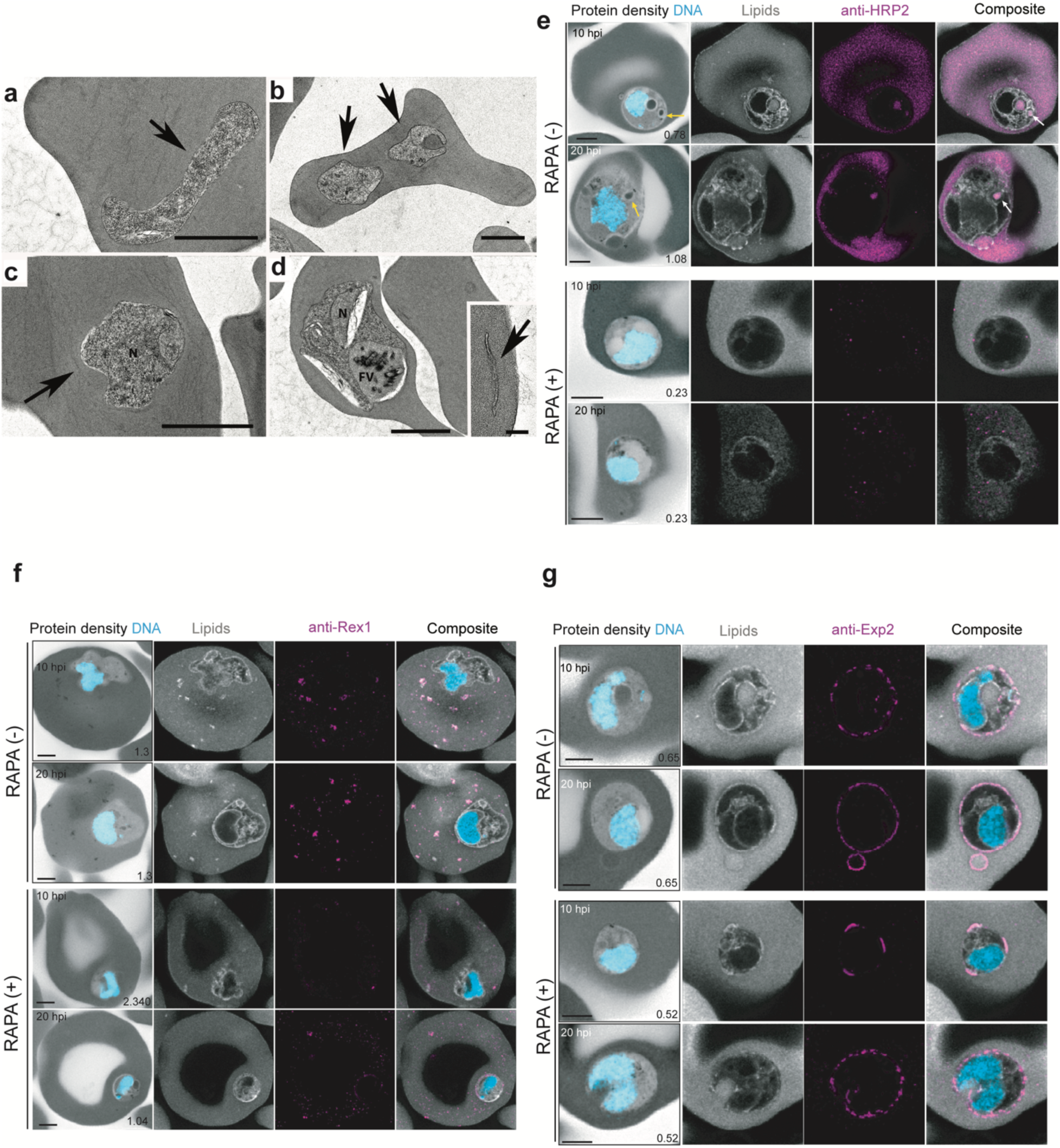
Truncation of *pfap2-hcr* affects key protein trafficking components. **a-d**, Electron micrographs of control (RAPA−) and treated (RAPA+) iRBCs at both 12 and 24 h.p.i. at cycle 1. **a,** Transmission electron micrograph of an RBC from RAPA+ iRBCs at 12 h.p.i. showing an early crescentic shaped parasite (arrow). **b**, RBC containing two early trophozoites from the RAPA+ iRBCs at 24 h.p.i. (arrows). **c**, Section through an infected RBC for the RAPA− iRBCs at 12 hrs showing an early trophozoite stage (arrow). **d**, Section through an infected RBC for the RAPA− iRBCs at 24 h.p.i. illustrating a late trophozoite with a central nucleus (N) and a large food vacuole (FV) containing hemozoin crystals. Bar is 1 µm. Insert. Detail of iRBC cytoplasm showing a Maurer’s cleft (arrow), Bar in insert 200nm. **e-g**, Control (RAPA−) and treated (RAPA+) PfAP2-HCR-3HA:*loxP* parasites were prepared through U-ExM, stained with N-hydroxysuccinimide (NHS) ester (grayscale), BODIPY FL (white), SYTOX (cyan), and antibodies against HRP2 (**e**), Rex1 (**f**), and Exp2 (**g**) (magenta), and imaged with Airyscan microscopy. In panel **e,** yellow arrows mark the cytostome and white arrows mark the presence of HRP2 signal (host cytoplasm) within the invagination of the cytostome. Images are maximum-intensity projections; the number indicates the Z-axis thickness of the projection in µm. Scale bars = 5 µm.

Notably, PfAP2-HCR directly bound and transcriptionally regulated two essential parasitophorous vacuole membrane (PVM) proteins, Exp1^50^ and Exp2^8,51^, the latter being a core constituent of the PTEX translocon (**Fig. 3e,f**; **Supplementary Data 1 and 2**). In addition, PfAP2-HCR directly controlled the expression of numerous other key exported proteins, including *hrp2*, *rex* family members, *resa*, and multiple *etramps*, implicating it as a central regulator of the export machinery. To assess the consequences of *pfap2-hcr* truncation on the expression and subcellular localization of these targets, we performed U-ExM using antibodies against selected exported proteins at 10 and 20 h.p.i. in cycle 1, following rapamycin treatment at 5 h.p.i. in cycle 0. We examined the status of HRP2, a soluble PEXEL-positive exported protein, and Rex1, a PEXEL-negative Maurer’s cleft (MC)–resident protein. As expected, HRP2 signal was readily detected throughout the infected red blood cell (iRBC) cytoplasm in control parasites, whereas *Δpfap2-hcr* parasites exhibited markedly reduced or absent HRP2 signal (**Fig. 5e**). In contrast, Rex1 localized to punctate structures within the iRBC cytoplasm in control parasites, consistent with normal Maurer’s cleft formation; however, Rex1 signal was almost completely lost in *Δpfap2-hcr* parasites, indicating a failure of expression and/or MC biogenesis (**Fig. 5f**). These observations are concordant with transmission electron microscopy analyses, which revealed an absence of Maurer’s clefts in iRBCs infected with *Δpfap2-hcr* parasites.

In *Δpfap2-hcr* parasites, we further assessed the expression and subcellular distribution of Exp2 as well as two additional core components of the PTEX translocon, HSP101 and PTEX150 in iRBCs. In control parasites, Exp2 displayed a continuous and uniform localization along the PVM. In contrast, *Δpfap2-hcr* parasites exhibited a discontinuous and patchy staining at the parasite periphery at both time points examined, consistent with reduced expression and/or impaired targeting of Exp2 to the PVM (**Fig. 5g**). By comparison, the localization of HSP101 and PTEX150 was largely unaffected, with only minor or no detectable differences between control and *Δpfap2-hcr* parasites (**Extended Data Fig. 10**). These observations indicate a selective disruption in Exp2 abundance and spatial organization, rather than a uniform loss of PTEX components, and suggest that PfAP2-HCR is required to ensure proper assembly or distribution of the export translocon at the PVM. Collectively, these data demonstrate that truncation of *pfap2-hcr* leads to profound defects in the expression and organization of essential exported proteins and PVM-associated components, resulting in a severe impairment of host cell remodeling during asexual blood-stage development.

## Discussion

In this study, we identified PfAP2-HCR as an essential transcriptional regulator that orchestrates the expression of exported proteins required for host cell remodeling during the ring stage of *Plasmodium falciparum* development. Although PfAP2-HCR was initially identified as one of the downstream targets of the circadian-expressed regulator PfAP2-P ^31^, our results revealed that it forms a key node within a broader transcriptional network governing early intraerythrocytic development. Conditional truncation of *pfap2-hcr* led to a striking arrest immediately after erythrocyte invasion, demonstrating that PfAP2-HCR is indispensable specifically during the ring stage maturation period. This phenotype contrasts with the initially normal egress and invasion of *Δpfap2-hcr* parasites, underscoring that PfAP2-HCR is not required for merozoite formation or invasion but instead functions immediately thereafter to initiate the host cell remodeling program, which is critical for further intraerythrocytic development.

The RNA-seq data demonstrated that PfAP2-HCR predominantly acts as a transcriptional activator. Loss of PfAP2-HCR resulted in substantial downregulation of genes enriched for Maurer’s cleft, host cell surface, and parasitophorous vacuole membrane components, which are structures central to protein export and erythrocyte modification. Notably, over 23% of the entire *P. falciparum* exportome ^15^ was downregulated, indicating that PfAP2-HCR regulates a large portion of exported protein genes. These effects were particularly pronounced in Cluster 3, which comprised genes that peaked during the late schizont and early ring stages and encoded key determinants of host cell remodeling including *rex*, *resa*, *etramp*, *sbp1*, *fikk* kinases, and *emp1-trafficking proteins*. The enrichment of PfAP2-HCR targets within this expression cluster strongly supports a model in which PfAP2-HCR primes the nascent ring stage parasite to remodel its intracellular niche immediately after its invasion.

ChIP–seq analysis confirmed that PfAP2-HCR directly binds to the putative promoters of 51 exported protein–encoding genes, including the essential PVM-resident proteins Exp1^52^ and a critical PTEX translocon component Exp2 ^8,51^.

U-ExM–based phenotyping was critical for linking these PfAP2-HCR–dependent transcriptional changes to the spatial disorganization and functional collapse of the export machinery within infected erythrocytes. Consistent with this, *Δpfap2-hcr* parasites showed a drastic reduction in exp2 expression. U-ExM analyses of soluble and Maurer’s cleft–resident exported proteins further reveal how PfAP2-HCR loss translates into a collapse of the host cell remodeling machinery. HRP2, a soluble PEXEL-positive protein, shows robust signal throughout the infected erythrocyte cytoplasm in control parasites, whereas HRP2 signal is markedly reduced or absent in *Δpfap2-hcr* parasites, consistent with impaired expression and/or export. Likewise, the PEXEL-negative Maurer’s cleft marker Rex1 localizes to discrete puncta in control parasites but is almost completely undetectable in *Δpfap2-hcr* parasites, indicating a failure of Maurer’s cleft biogenesis that aligns with the absence of clefts by transmission electron microscopy. Together, the U-ExM and electron microscopic data show that PfAP2-HCR-dependent transcription is required not only to establish the PVM export gateway but also to build the downstream trafficking and sorting platforms that distribute exported cargoes within the infected red blood cell.

Beyond the exportome, PfAP2-HCR binds to the putative promoter regions of multiple other ApiAP2 transcription factors, including *ap2-g*, *ap2-exp*, *ap2-o*, and *ap2-l*, and associates with chromatin-remodeling proteins such as PfMORC^38,48^, PfISWI^39^, and PfSET10^40^. These interactions place PfAP2-HCR within a combinatorial regulatory network similar to other ApiAP2 hubs such as PfAP2-P and PfAP2-I ^29,31^ and suggest that its regulatory activity is embedded within a higher-order chromatin architecture.

An unexpected observation was that PfAP2-HCR binds to the putative promoters of subtelomeric *var* genes and that Δ*pfap2-hcr* parasites show widespread downregulation of multiple *var* loci. Because *var* genes are transcriptionally silent during the stages used for RNA-seq and ChIP-seq, and because directly bound *var* promoters do not correspond to downregulated *var* genes, these effects are likely indirect, potentially reflecting perturbation of heterochromatic organization mediated through PfAP2-P, PfMORC- or PfISWI-associated complexes.

Our findings establish PfAP2-HCR as a key regulator of the early ring stage remodeling program and highlights its broader implications for parasite biology. PfAP2-HCR is highly expressed in multiple life-cycle stages, including sexually committed schizonts and rings, ookinetes, and sporozoites (**Extended Data Fig. 2a)**, suggesting potential functions beyond asexual blood-stage development. Gametocytes extensively remodel erythrocytes to support sequestration and maturation ^3^, while liver-stage parasites reorganize hepatocyte architecture during intracellular expansion^53–58^.

Consistent with its expression in sexual-stage development, conditional truncation of *pfap2-2hcr* impaired gametocytogenesis, resulting in developmental arrest and the accumulation of pyknotic or deformed gametocytes. This phenotype suggests that PfAP2-HCR regulates exported proteins required for host-cell remodeling during early gametocytogenesis. Notably, PTEX components required for protein export are expressed in stage I/II gametocytes but are degraded during later gametocyte maturation^9^, indicating that export is most active during the earliest stages of gametocytogenesis^59^. This early requirement for export provides a plausible explanation for the high proportion of sexually committed rings that became pyknotic or morphologically abnormal following *pfap2-hcr* truncation.

Further supporting this model, several *P. falciparum* gametocyte-exported protein genes were downregulated in *Δpfap2-hcr* parasites, including *pfgexp5*^60^, *pfgexp12*, *pfgexp21* and the gametocyte erythrocyte cytosolic protein *pfgeco*^61^ (Supplementary Data 1). Given that PfAP2-HCR regulates numerous essential exportome genes and is highly expressed in ookinetes, sporozoites and potentially liver-stage parasites, defining its function in these stages may reveal additional stage-specific role, exported proteins and remodeling pathways. This is particularly important for liver-stage development, where parasite-mediated host-cell remodeling remains comparatively poorly understood.

Overall, our work identifies PfAP2-HCR as a central transcriptional regulator of host cell remodeling required for ring stage maturation and gametocyte development (**Fig. 6**). The broad expression of PfAP2-HCR in merozoites, rings, ookinetes, and sporozoites, coupled with its regulation of numerous essential exported proteins, suggests that analogous remodeling modules may operate in non-erythrocytic stage like liver stage. The U-ExM framework established here provides a powerful approach to dissect how PfAP2-HCR-controlled exportome sculpt distinct host-cell environments, particularly in liver stages where remodeling remains poorly defined. Future work combining stage-specific perturbation of PfAP2-HCR with super-resolution imaging and functional assays in gametocytes and liver stages should reveal whether a conserved PfAP2-HCR–driven remodeling logic underlies multiple transitions in the parasite life cycle and may identify exported proteins that constitute attractive targets for multistage interventions.

**Fig. 6.**
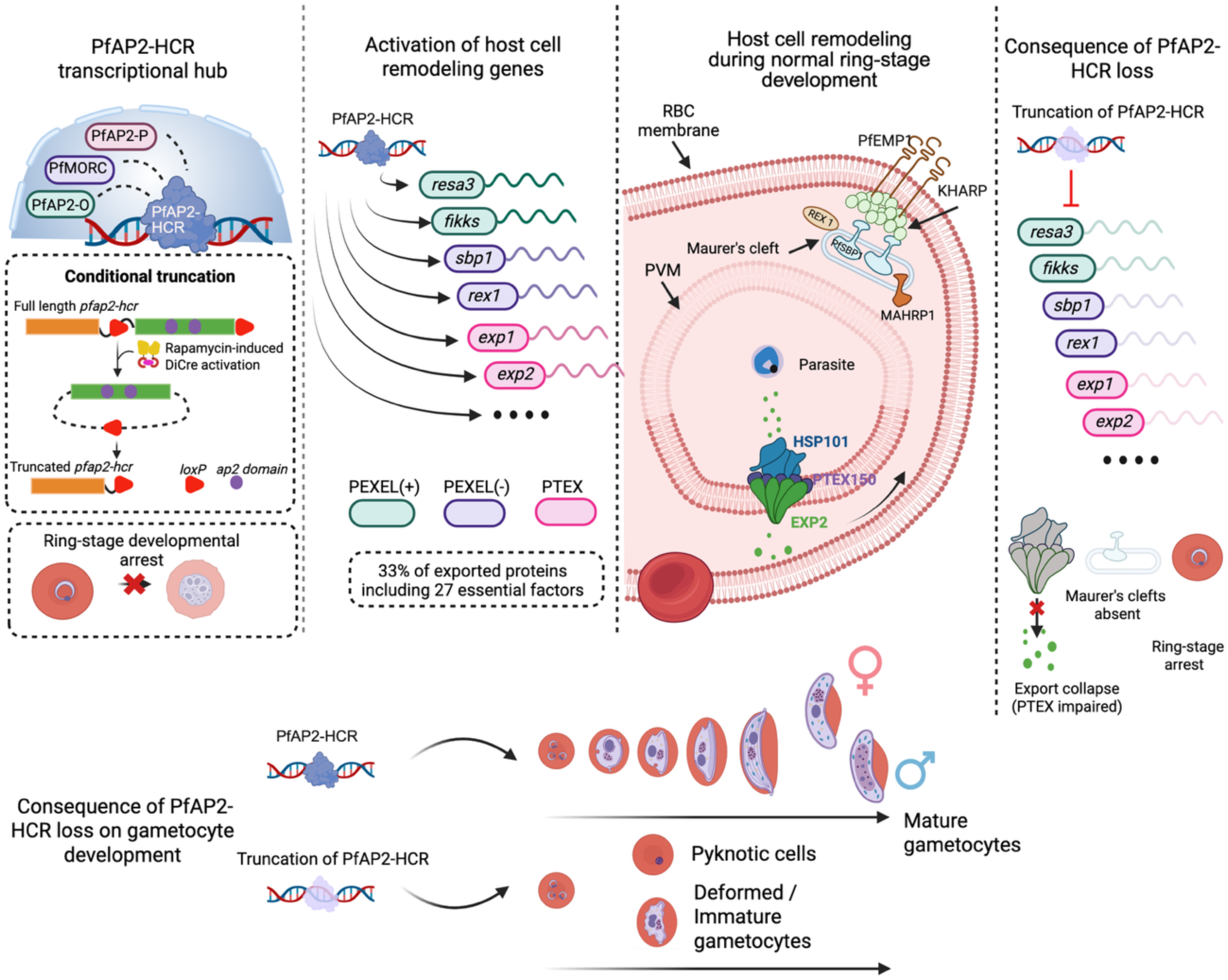
PfAP2-HCR coordinates expression of exported proteins required for host cell remodeling during blood-stage development. Schematic model summarizing the proposed role of PfAP2-HCR as an essential transcriptional regulator of host cell remodeling in *Plasmodium falciparum*. PfAP2-HCR functions within a chromatin-associated regulatory network that includes PfAP2-P, PfMORC and PfAP2-O, and promotes the expression of exported protein required for host-cell remodeling. Conditional truncation of *pfap2-hcr* is induced by rapamycin-mediated DiCre activation, resulting in l*oxP*-dependent excision of the gene encoding C-terminal region of the protein including the AP2 DNA-binding domains. PfAP2-HCR truncation results in impaired export machinery, absence or collapse of Maurer’s clefts, defective host cell remodeling and ring stage developmental arrest. PfAP2-HCR truncation further affects gametocyte development and maturation. The model highlights PfAP2-HCR as a central transcriptional hub required for parasite-induced host cell remodeling during asexual ring stage and gametocyte development to support blood stage development.

## Materials and Methods

### Parasite culture, maintenance and synchronization

The DiCre-expressing *Plasmodium falciparum* clone II-3 ^46^ was used for all conditional gene truncation experiments. Parasites were maintained in human AB⁺ erythrocytes at 3% haematocrit in RPMI 1640 supplemented with Albumax II (Invitrogen) and 2 mM L-glutamine, and cultured at 37 °C under standard gas conditions. Parasite synchronization was performed either by 5% sorbitol treatment or by Percoll-based enrichment of mature schizonts. For Percoll synchronization, late schizonts were purified using 63% Percoll, allowed to invade fresh erythrocytes for 1–2 h, followed by sorbitol treatment to obtain tightly synchronized ring stage parasites.

### Transfection of *P. falciparum*

For genome editing, ∼10⁸ Percoll-enriched mature schizonts were resuspended in 100 µl P3 Primary Cell Solution (Lonza) containing 60 µg linearized repair plasmid DNA (repair plasmid 1 or 2, in separate experiments) and 20 µg pDC2-derived guide RNA plasmid. Transfections were performed using an Amaxa 4D-Nucleofector (Lonza) with program FP158, as described previously ^62^. Drug selection was initiated ∼20 h post-transfection using 5 nM WR99210 (a kind gift from Jacobus Pharmaceuticals) for four days. Once stable growth was established, parasites were subjected to negative selection with 1 µM 5-fluorocytosine (Ancotil, MEDA) for four days. Clonal lines of PfAP2-HCR-3HA:loxP parasites were obtained by limiting dilution. Conditional excision mediated by DiCre recombinase was induced by treatment with 10 nM rapamycin.

### Gametocyte culture and maintenance

Sorbitol synchronized, high parasitemia PfAP2-HCR-II3 parasite cultures were used for gametocytogenesis induction. ITSx supplement (1X) was used as an inducing agent^63^ along with 50 mM N-acetyl glucosamine (NAG) to restrict asexual parasite growth. At ring stage, parasites were induced for gametocytogenesis by supplementing growth media with 1X ITSx supplement and 50 mM NAG (day 0). Till day 4, media was changed every day, later, every alternate day till day 10. On day 10, gametocytes/pyknotic parasites were enriched using percoll (63% percoll). *Pfap2-hcr* truncation was carried out by addition of 10 mM rapamycin under two different conditions: 1) HCR truncation followed by gametocytogenesis induction (addition of DMSO/RAPA on day -1 (late trophozoite stage) and addition 1X ITSx + 50 mM NAG on Day 0) and 2) HCR truncation along with gametocytogenesis induction (addition of DMSO/RAPA + 1X ITSx + 50 mM NAG on Day 0). 48 hrs post RAPA treatment, parasites were sampled for HCR truncation by excision PCR using Takara CloneAmp master mix and HCR specific primers. Smears were made and stained by Giemsa staining. Images were acquired using Zeiss Axiocam 512 microscope using 100x oil immersion objective. Gametocytes (Mature/immature/deformed) or pyknotic parasites were counted manually from percoll enriched day 10 Giemsa-stained images.

### Strategy for conditional truncation of *pfap2-hcr*

To delete the genomic region encoding both AP2 domains, we generated a two-step Cas9-mediated “floxed” and HA-tagged allele of *pfap2-hcr* in the II-3 DiCre background. Step 1: The C terminus of PfAP2-HCR was tagged with 3×HA, and a loxP site was inserted immediately downstream. The repair plasmid (GenScript, USA) contained recodonized sequence corresponding to nucleotides 9865–10419 of *pfap2-hcr*, flanked by 400 bp 5′ and 3′ homology arms. Step 2: A second *loxP* site (*sera2*-*loxPint*) was introduced upstream of AP2 domain 1 using repair constructs containing recodonized sequences corresponding to nucleotides 4714–4896 and 400 bp flanking homology regions. All edits were performed using the pDC2-Cas9-U6-hDHFR plasmid expressing *S. pyogenes* Cas9. Guide RNAs were designed using Benchling. Sequences of all oligonucleotides and primers are provided in **Supplementary Data 6**.

### Growth assay

For growth assay, synchronous ring stage parasites at 0.1% parasitemia and 3% hematocrit were maintained in triplicate in 24-well plates. Fifty microliters of sample per well was collected at ∼24 h.p.i. in each replication cycle starting from cycle 0 until cycle 2 for both mock-treated control (DMSO) and RAPA-treated samples (3 replication cycles), fixed with 50μl of 0.2% glutaraldehyde in phosphate-buffered saline (PBS) and stored at 4°C for flow cytometry-based quantification. Fixed parasites were stained with SYBR Green (Thermo Fisher Scientific, 1:10 000 dilution) for 30 min at 37°C and parasitemia was determined using a BD LSR Fortessa flow cytometer (BD Biosciences).

### Nucleic acid extraction and polymerase chain reaction

For genomic DNA isolation, RBCs were pelleted and treated with 0.15% saponin in phosphate-buffered saline (PBS) for 10 min on ice, followed by two washes with PBS. DNA was extracted from the resulting parasite pellets using the DNeasy Blood and Tissue Kit (Qiagen) according to the manufacturer’s instructions. Diagnostic PCRs for confirmation of parasite clones were performed using CloneAmp HiFi polymerase (Takara).

### RNA extraction and strand-specific RNA-seq library preparation

Parasite cultures (0.5–2 ml, depending on developmental stage) were pelleted at 2,400 rpm for 3 min and lysed in 1 ml TRIzol (Sigma) by passing through 1 ml tips at least 10 times. Lysates were immediately stored at −80 °C until processing. Total RNA was extracted using the TRIzol reagent protocol (Life Technologies) according to the manufacturer’s instructions.

Strand-specific mRNA libraries were prepared using the TruSeq Stranded mRNA Sample Prep Kit LS (Illumina). For each sample, ∼500 ng of total RNA was used as input. Poly(A)⁺ RNA was enriched using oligo-dT–coated magnetic beads, followed by first-strand synthesis using random primers. Second-strand synthesis incorporated dUTP in place of dTTP to ensure strand specificity. After end repair and adaptor ligation, libraries were amplified by PCR for 15 cycles and sequenced on an Illumina HiSeq 4000 platform using 150-bp paired-end reads.

### RNA-seq data processing and analysis

Raw sequencing reads were assessed for quality using FASTQC v0.12.8, and Illumina adaptor sequences and low-quality bases were trimmed using Trimmomatic^64^. Processed reads were aligned to the *P. falciparum* 3D7 reference genome (PlasmoDB release 67; http://www.plasmodb.org using HISAT2 v2.2.1^65^ with the parameter --rna-strandness RF. Gene-level read counts were obtained using FeatureCounts^66^.

Reads were converted to counts per million (CPM), and genes with CPM <1 in all three biological replicates were excluded. Library size normalization was performed using the trimmed mean of M-values (TMM) method implemented in EdgeR v3.40.2^67^, and normalized counts were transformed using the *voom* function in the limma package^68^. Differential expression analysis was conducted using DESeq2 v1.38.3^69^. Genes with a Benjamini–Hochberg false discovery rate (FDR) <0.05 and absolute log₂ fold-change ≥1 or ≤−1 were classified as significantly upregulated or downregulated, respectively.

### Chromatin immunoprecipitation (ChIP)

ChIP was performed as previously described^31^ with modifications. Coumpund 2-treated parasites at very late segmented schizont stage with ∼5% parasitemia were pelleted at 900 × *g* for 4 min and washed once in PBS. Parasites were released by incubation with 25 ml of 0.15% saponin in PBS for 10 min on ice, followed by two washes with cold PBS. Cross-linking was performed by adding methanol-free formaldehyde to a final concentration of 1% and incubating at 37 °C for 10 min with intermittent mixing. The reaction was quenched by adding glycine to a final concentration of 0.125 M and incubating the mixture for an additional 5 min at 37 °C. Parasites were pelleted at 3,250 × *g* for 10 min at 4 °C, washed thrice in DPBS, snap-frozen in liquid nitrogen, and stored at −80 °C.

For ChIP, frozen cross-linked pellets were thawed on ice and resuspended in 1 ml nuclear extraction buffer (10 mM HEPES, 10 mM KCl, 0.1 mM EDTA, 0.1 mM EGTA, 1 mM DTT, and 1× EDTA-free protease inhibitor cocktail (Roche)). After 30 min on ice, NP-40 was added to a final concentration of 0.25%, and the parasites were lysed by passage through a 26½-gauge needle seven times. Nuclei were pelleted at 2,500 × *g* for 10 min at 4 °C.

Chromatin was sheared using a Covaris E220 sonicator for 14 min (5% duty cycle, 140 peak incident power, 200 cycles/burst) to obtain fragments of 200–600 bp. Insoluble debris was removed by centrifugation at 13,500 × *g* for 10 min at 4 °C, and 30 µL of sheared chromatin was reserved as input control and stored at −80 °C.

The remaining chromatin was diluted 1:1 in ChIP dilution buffer (30 mM Tris–HCl pH 8.0, 0.1% SDS, 3 mM EDTA, 300 mM NaCl, 1% Triton X-100, plus protease inhibitors) and incubated overnight at 4 °C with anti-HA antibody (Abcam ab9110). Immune complexes were captured using protein A–coupled magnetic Dynabeads (Invitrogen 10002D), washed sequentially with low-salt wash buffer, high-salt wash buffer (4 °C), and TE buffer (room temperature).

Chromatin was eluted in 1% SDS, 0.1 M NaHCO₃ at 45 °C for 30 min with shaking. Input and immunoprecipitated samples were reverse cross-linked overnight at 45 °C after adding NaCl to 0.5 M. Samples were treated with RNase A for 30 min at 37 °C and then incubated with proteinase K (66 µg ml⁻¹) for 2 h at 45 °C. DNA was purified using the ChIP DNA Clean & Concentrator kit (Zymo Research D5205).

### ChIP–seq library preparation, processing and analysis

ChIP–seq libraries were prepared using the NEBNext Ultra II DNA Library Prep Kit (NEB) following the manufacturer’s protocol up to the adapter-ligation step, with adapters diluted 1:20. Adapter-ligated DNA was purified using AMPure XP beads, and libraries were amplified for eight PCR following the manufacturer’s protocol. Amplified libraries were purified, size-selected for ∼350 bp inserts using AMPure XP beads, and sequenced on the Illumina HiSeqX platform using 150 bp paired-end reads.

Low-quality bases and Illumina adapters were removed using Trimmomatic v0.39^64^, and trimmed reads were mapped to the *P. falciparum* 3D7 genome (PlasmoDB v3, release 67) using HISAT2 v2.2.1^65^. Duplicate reads were removed using *Samtools* v1. (markdup)^70^. GC bias was corrected using deepTools v 3.3.1 (*correctGCBias*) ^71^. Coverage tracks for 16 h.p.i. and 40 h.p.i. ChIP–seq experiments were generated using deepTools v 3.3.1 *bamCompare* with base-pair resolution (-bs 1) and normalization to RPKM (--normalizeUsing RPKM). The input signal was subtracted from the ChIP coverage using the --operation subtract option. Tracks were visualized using IGV v2.4.16^72^.

ChIP peaks were called using MACS3 v3.0.1^73^ with a q-value threshold of <0.05, comparing input vs. ChIP. Peak calling was performed using -nomodel, an extension size of 200 bp (-extsize 200), and a genome size of 23,332,839 bp. Reproducible peaks across biological replicates were identified using *bedtools* v2.30.0 *intersect* (-f 0.30 -r)^74^ with at least 30% of the genomic region shared by each peak from the replicates. Peaks with scores >50 were retained for downstream analysis. The score was determined by the *bedtools* v2.30.0 *intersect* function and a cutoff score of 50 was determined empirically. Annotation to the nearest downstream gene was performed using HOMER’s (v 5.1) *annotatePeaks.pl* ^75^.

### PfAP2-HCR motif identification

Sequences corresponding to reproducible ChIP–seq peaks shared between biological replicates were retrieved from the *P. falciparum* 3D7 genome using *bedtools* v2.29.0 (*getfasta*). Enriched DNA motifs within these peak regions were identified using the DREME algorithm^76^. De novo motifs were subsequently compared to previously characterized ApiAP2-binding motifs using Tomtom^77^ and in silico motif databases^78^.

### Transmission Electron Microscopy

The cells were fixed with 2.5% glutaraldehyde in 0.1M Sodium Cacodylate buffer pH 7.4 at 4°C (Electron Microscopy Science). Within 24 h the samples were rinsed three times with fresh 0.1 M cacodylate buffer pH 7.4 (Electron Microscopy Science) at room temperature for 5 min. Cells were post-fixed using 1% osmium tetroxide (Electron Microscopy science) in 1.5% Potassium ferrocyanide for 1h at room temperature followed by 2% osmium tetroxide in water for 30 min at room temperature and rinsed three times with water for 5 min at room temperature after each incubation. The preparations were incubated with 1% Uranyl acetate at 4°C for 12h.

Subsequently, the cells were dehydrated in serial ethanol dilutions 50%, 70%, 80%, 90%, 100% followed by 100% acetone and infiltrated in 3:1, 1:1 and 1:3 acetone/Epon resin (Electron Microscopy Science) mixtures. Finally, the samples were embedded in pure Epon resin and cured for 72h at 60°C.

Ultrathin sections of 90 nm were prepared using UC6 (Leica Microsystems) and mounted on copper grids and eventually stained with 3% lead citrate (Electron Microscopy Science) for 4 min. Micrographs were acquired using Talos F200X transmission electron microscope (Thermofisher Scientific) operating at 200 kV.

### Ultrastructure Expansion Microscopy (U-ExM)

Ultrastructure Expansion Microscopy (U-ExM) was performed as previously described^79^, with minor adaptations for *Plasmodium falciparum* blood-stage parasites, based on protocols established in the laboratory for high-resolution imaging of parasite subcellular architecture^80,81^.

### Parasite preparation and fixation

Synchronized parasites at 6 h.p.i. in growth cycle 0 were treated with either DMSO or RAPA, and parasites at 10 and 20 h.p.i. in cycle 1 were fixed with pre-warmed 4% (v/v) paraformaldehyde (PFA) at 37 °C for 20 minutes. Fixed cells were washed in PBS, stored, and shipped to Absalon at 4 °C for U-ExM processing, immunostaining, and imaging. Fixed iRBCs were first settled on 12 mm poly-D-lysine–coated coverslips and processed for ultrastructure expansion microscopy (U-ExM). Briefly, coverslips were coated with poly-D-lysine for 1 hour at 37 °C, washed twice with MilliQ water, and placed in a 24-well plate. Parasite cultures adjusted to approximately 1% hematocrit were added to each well, and iRBCs were allowed to settle for 30 minutes at 37 °C before the supernatant was removed. Cells were fixed by gently adding 4% (v/v) PFA in 1× PBS along the side of the well and incubating for 20 minutes at 37 °C. Coverslips were washed once with 1× PBS and incubated in 1.4% (v/v) formaldehyde/2% (v/v) acrylamide (FA, 36.5-38%, F8775-25mL, SIGMA /AA 40%, A4058, SIGMA) in PBS overnight at 37 °C.

After PBS washes, samples were embedded in a monomer solution containing sodium acrylate (97-99%, 408220, SIGMA), acrylamide (40%, A4058, SIGMA), and N,N′-methylenebisacrylamide (2%, M1533, SIGMA), supplemented with ammonium persulfate (APS) and TEMED. Polymerization was conducted at 37 °C for 30 minutes. Gel denaturation, expansion, and immunostaining were performed as previously described. Expanded gels were blocked with 3% BSA in PBS for 30 minutes at room temperature with shaking.

Primary antibodies were diluted in blocking buffer and applied to gels for an overnight incubation at room temperature with gentle agitation, followed by three washes with 0.1% Triton-PBS. Gels were then incubated with mouse or rabbit Alexa Fluor 555–conjugated secondary antibodies (1/500, #A21422, #A32732 ThermoFisher), NHS-ester-405 (1/200, #A30000 ThermoFisher), and Sytox Deep Red (1/1000, #S11380 ThermoFisher) for 48 hours at 4 °C. The following antibodies were used: anti-Exp2 (1/500, rabbit), anti-HSP101 (1/250, rabbit,), anti-PTEX150 (1/250, rabbit)-all three antibodies generously gifted by Dr. Paul Gilson^82^. anti-HRP2 (1/200, mouse, #MA5-18245, ThermoFisher), anti-Rex1 (1/1000, rabbit, generously gifted by Dr. Tobias Spielman)^52^, and anti-HA (1/400, rabbit, #AF291 from ABCD). For cellular context, gels were additionally stained with BODIPY™ FL C5-Ceramide (1/500, #D3521, ThermoFisher) in 2% Propyl Gallate solution at room temperature overnight with shaking to label membranes.

Stained gels were imaged using a Zeiss LSM900 AxioObserver equipped with an Airyscan 2 detector and a 63× Plan-Apochromat 1.4 NA objective. Z-stacks were captured with Nyquist sampling. Image acquisition and processing (including background subtraction, intensity adjustment, and overlay creation) were carried out in ZEN Blue (v3.5, Zeiss), and final figure assembly and maximum-intensity projections were prepared in Zen Blue. U-ExM experiments were repeated across independent biological replicates, and representative images are shown.

### Immunoprecipitation of PfAP2-HCR and mass spectrometric identification of associated proteins

To identify PfAP2-HCR–associated proteins, the immunoprecipitated complexes were subjected to on-bead trypsin digestion. Following the final Tris–EDTA washes, the beads were washed twice with exchange buffer (100 mM NaCl, 50 mM Tris pH 7.5) for 10 min at 4 °C. The beads were then resuspended in 100 µL of 100 mM triethylammonium bicarbonate, and bound proteins were reduced with 1 mM dithiothreitol (DTT) at 37 °C for 30 min with shaking (750 rpm). The samples were equilibrated to room temperature and alkylated in the dark with 3 mM iodoacetamide for 45 min, followed by quenching with 3 mM DTT for 10 min. Proteins were digested overnight at 37 °C with 2.5 µg of trypsin (Promega) on a thermomixer (1,000 rpm).

The digested peptides were separated from the bead material, acidified with trifluoroacetic acid (TFA) to a final concentration of 2%, and desalted using Sep-Pak C18 cartridges (Waters). The columns were conditioned with methanol (2 × 1 ml) and equilibrated with 0.1% TFA (2 × 1 ml) prior to loading. The bound peptides were washed with 0.1% trifluoroacetic acid (TFA) (2 × 1 ml) and eluted twice with 300 µL of 75% acetonitrile/0.1% TFA. The eluates were dried using SpeedVac and stored at −80 °C until LC–MS analysis.

### LC–MS analysis of peptides

LC-MS/MS analysis was carried out on the Orbitrap Fusion Lumos mass spectrometer (ThermoFisher Scientific) coupled to a Dionex UltiMate 3000 RSLC nano HPLC system (Thermo Scientific). The peptide samples were loaded onto an Acclaim PepMap^TM^ 100 C18 trap column (100 µm x 2 cm, 100 A; Thermo Scientific) and resolved on a PepMap RSLC C18 analytical column (2µm, 100A, 75 µm x 50 cm; Thermo Scientific) at a flow rate of 300 nL/min over a 75 minutes gradient. Water with 0.1% FA (solvent A) and 95% ACN with 0.1% FA (solvent B) were used as mobile phases. The peptides were eluted from the column over a 55 minutes multistep gradient using 2.1% to 31.6% of solvent B followed by 5 minutes gradient to 90% B. The column was then maintained at 90% B for another 3 minutes to ensure elution of all the peptides before final equilibration with 2.1% B for 10 minutes.

The spray was initiated by applying 2 kV to the EASY-Spray emitter and the data were acquired under the control of Xcalibur software in a data-dependent mode using top speed and 3 s duration per cycle. The Orbitrap was operated in positive ionization mode, using the lock mass option (reference ion at m/z 445.120025) to ensure more accurate mass measurements in FTMS mode. The survey scan was recorded covering the *m*/*z* ranges from 400 and 1600 Th in profile mode with a resolution set to 120,000. The automatic gain control (AGC) target was set to standard and the maximum injection time was set to 50 ms. The most intense ions with charge state 2+ to 6+ were selected for fragmentation using HCD with a fixed collision energy mode. The HCD collision energy was set to 30% and the precursor ions were isolated using a 1.6 Th window. The AGC target was set to standard with a maximum injection time mode set to dynamic. The spectra were acquired using a 15,000 resolution

### Identification, quantification and statistical analysis of LC–MS data

Raw Q-Exactive HF data files were converted to *.mgf* format using ProteoWizard MSConvert (64-bit). Peptide identification was performed using Mascot v2.4 against the *Plasmodium falciparum* 3D7 protein database (PlasmoDB Release 51). Searches were performed with trypsin specificity allowing one missed cleavage, fixed carbamidomethylation of lysines, and variable modifications including deamidation (N/Q) and oxidation (M). The precursor and fragment mass tolerances were set to 0.6 Da.

## Acknowledgments

A.P. acknowledges support from King Abdullah University of Science and Technology (KAUST) via a faculty baseline fund (BAS/1/1020-01-01) and the KAUST Office of Sponsored Research—Competitive Research Grants (CRG-2020, CRG-2022) and the Opportunity Fund Program (OFP-2023)—under awards URF/1/4416-01-01, URF/1/5026-01-01 and URF/1/5501-01-01. S.A. is supported by an award from the Indiana University School of Medicine (S.A., BRG award no.2286272). We thank the staff of the Bioscience, Proteomics (under Analytical), and Imaging and Characterization Core Laboratories at KAUST for bulk RNA-seq library sequencing, flow cytometry (FACS), electron microscopy, and mass spectrometry–based experiments, and all members of the Pathogen Genomics Laboratory at KAUST for assistance during the experiments. Genomic data and bioinformatics tools were accessed through PlasmoDB (https://plasmodb.org), part of the VEuPathDB resource. We thank the PlasmoDB team for maintaining and continuously updating this essential resource for malaria parasite research.

## Data Availability

The datasets generated in this study are available in the following database. RNA-seq data: NCBI Bio Project accession no GSE322607; AP2-HCR ChIP– seq data: NCBI BioProject accession no. GSE322940; proteomics data: Pride accession number no. PXD075692. Bulk RNA-seq and ChIP–seq datasets have been added under the super series GSE322941. Source data are provided with this paper.

## Author Contributions

Conceptualization: A.K.S. and A.P.; methodology: A.K.S.; investigation: A.K.S., D.V., S.M., R.A.S., R.P.S., R.S., D.L., M.A., A.F.K., A.D., S.A.; analysis: A.K.S., S.A., D.J.P., R.S.; writing—original draft: A.K.S.; writing—review and editing: A.K.S., S. A. and A.P.; resources: S.A., and A.P.; funding acquisition: S.A. and A.P.; supervision: A.K.S., and A.P. All the authors have read and approved the manuscript.

## Corresponding Authors

Correspondence to Amit Kumar Subudhi and Arnab Pain

## Competing Interests

The authors declare no competing interests

## Supplementary Data Legends

**Supplementary Data 1 List of differentially expressed genes at schizont and ring stage after excision of *pfap2-hcr*.**

**Supplementary Data 2 List of genes encoding exported proteins clustered based on expression pattern during IDC.**

**Supplementary Data 3 ChIP–seq peaks identified in compound 2 treated PfAP2-HCR 3XHA tagged parasite at late schizont stage.**

**Supplementary Data 4 ChIP–seq peaks identified at 20, 30 and 40 h.p.i.**

**Supplementary Data 5 PfAP2-HCR associated proteins identified during late schizont stage using immunoprecipitation and mass spectrometry.**

**Supplementary Data 6 Oligos used in this study.**

---

**Extended Data Fig. 1.**
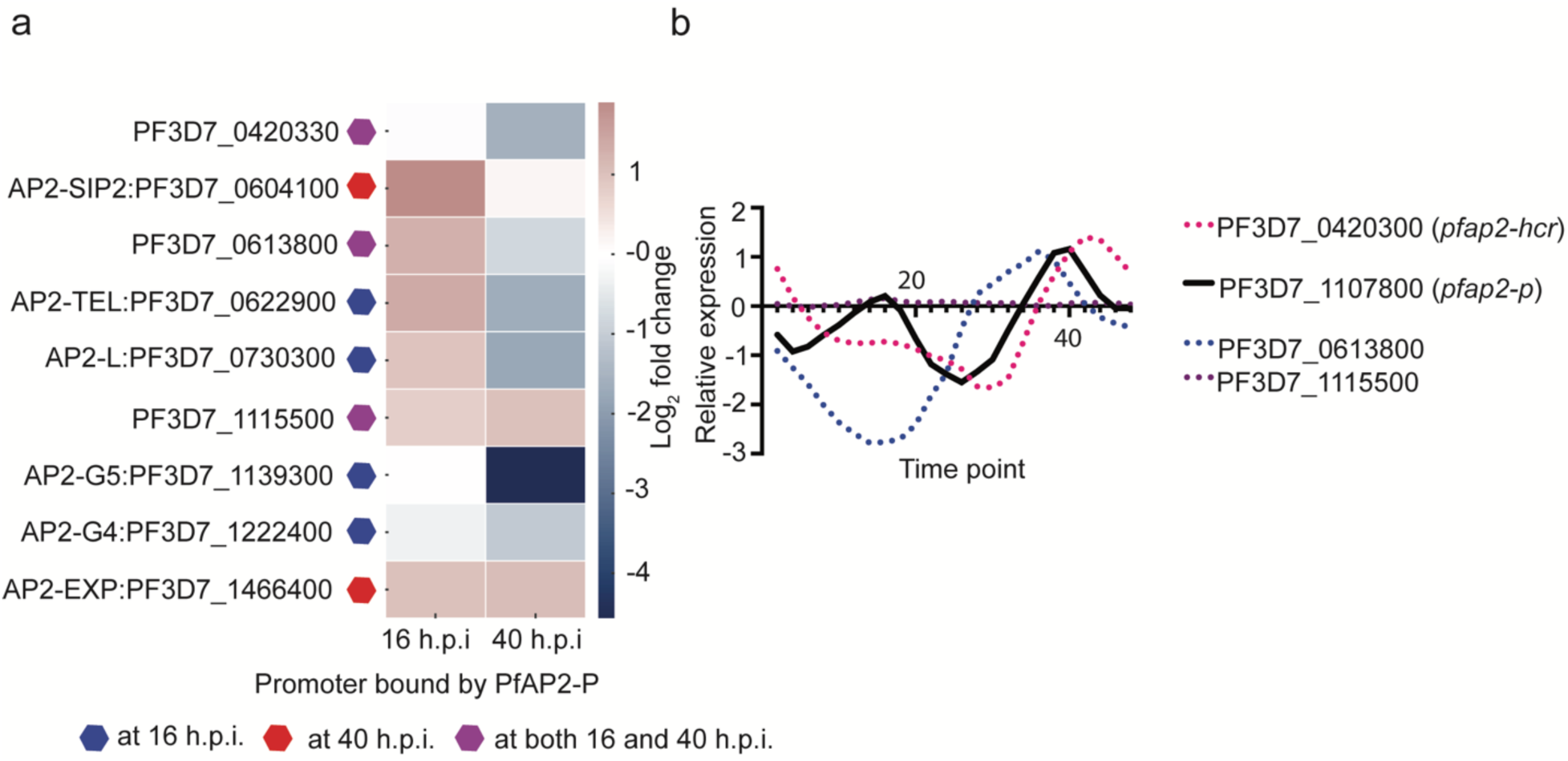
PfAP2-HCR is regulated by PfAP2-P. **a**, Heatmap showing the expression profiles of ApiAP2 transcription factors directly regulated by PfAP2-P during schizont-stage development, as reported previously^31^. **b**, Line graph showing the expression pattern of *pfap2-hcr* and other uncharacterized *pfap2s* compared to its upstream regulator *pfap2-p* across the intraerythrocytic developmental cycle (IDC)

**Extended Data Fig. 2.**
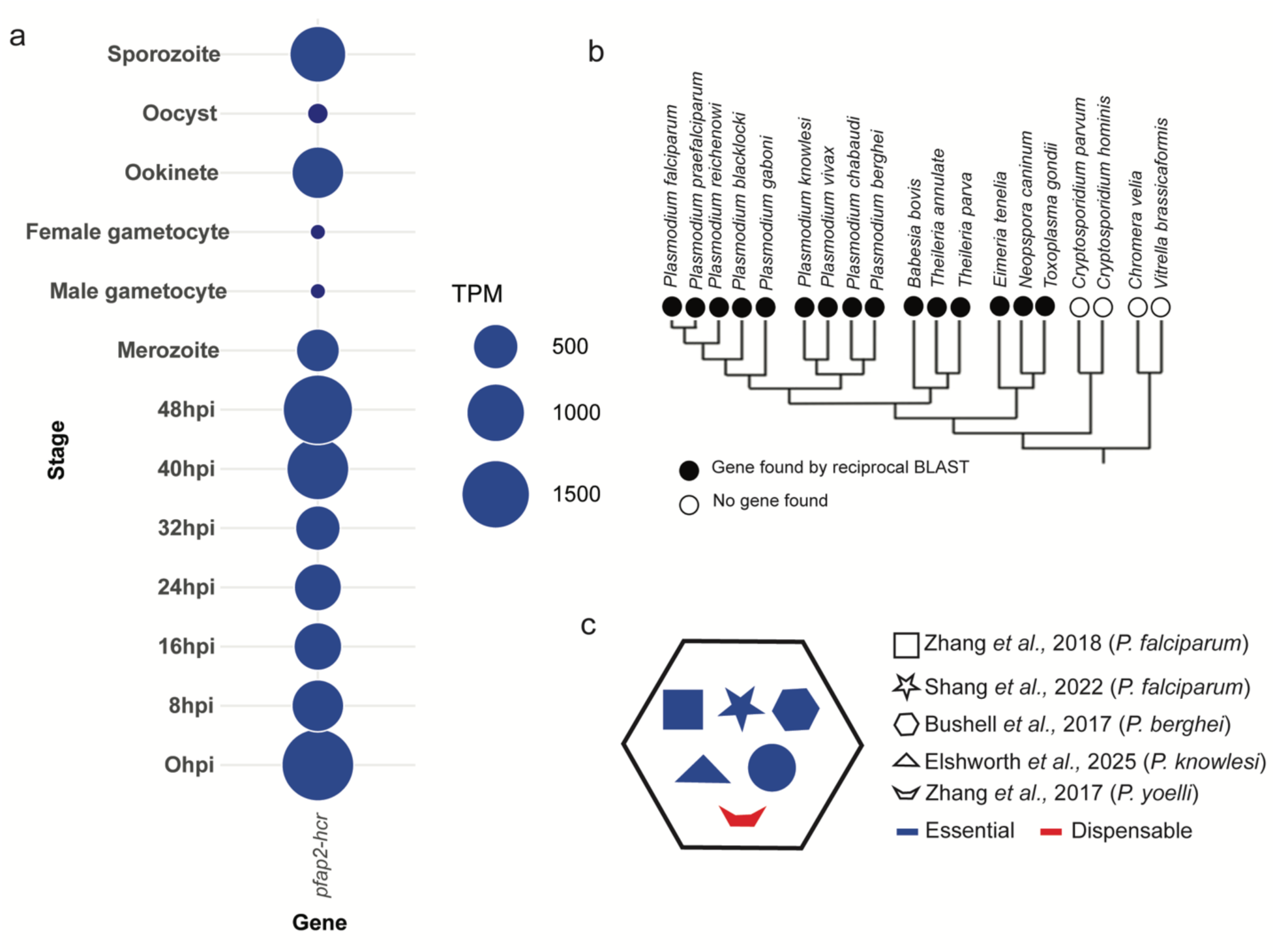
Basic features of PfAP2-HCR. **a,** Expression profile of *pfap2-hcr* throughout the *P. falciparum* life cycle (expression data collected from PlasmoDB). **b,** Phylogenetic distribution of PfAP2-HCR orthologs across the phylum *Apicomplexa.* **c,** Essentiality status of *pfap2-hcr* based on previous high-throughput genome-wide gene knockout studies.

**Extended Data Fig. 3.**
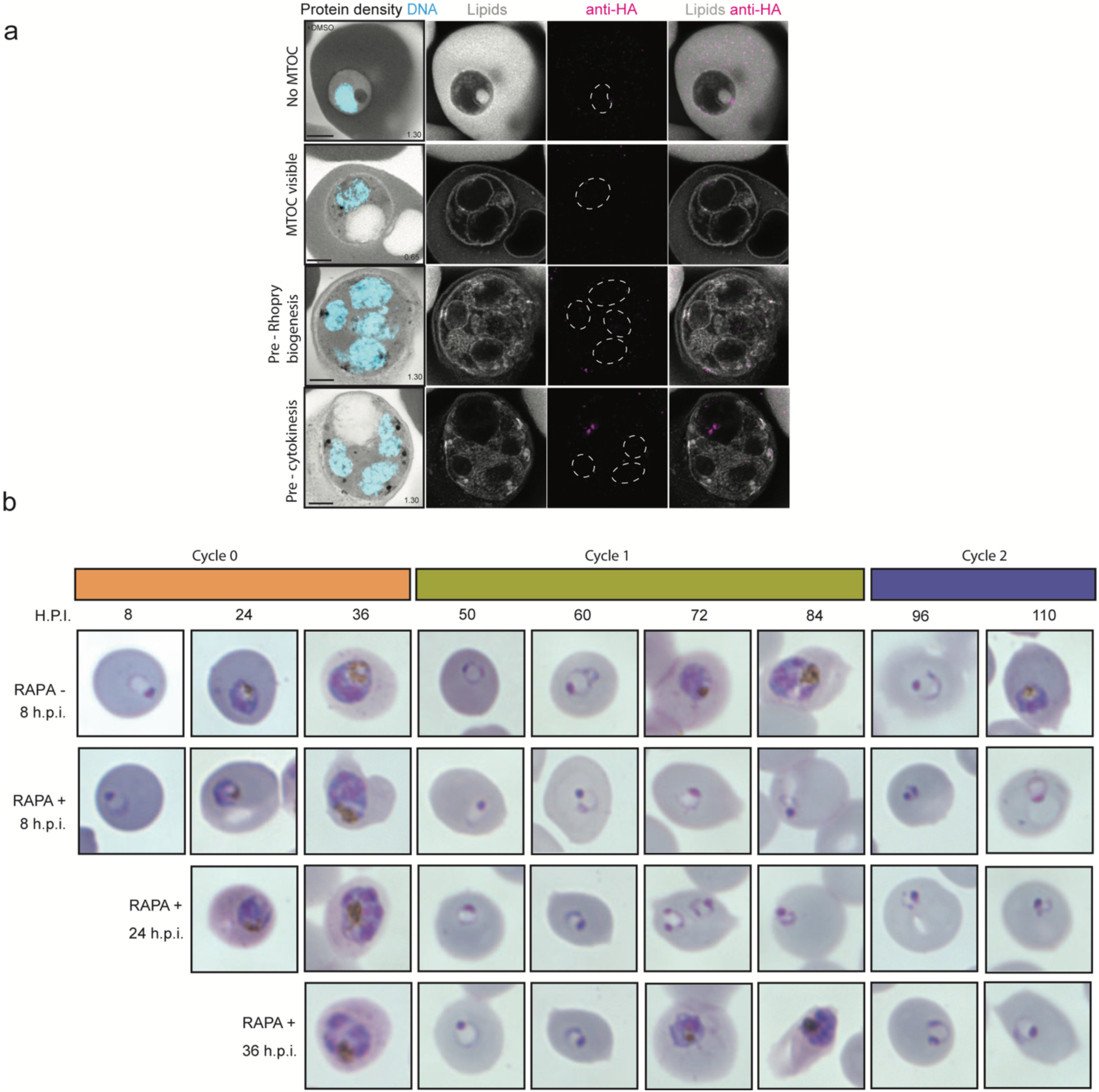
PfAP2-HCR is specifically essential during ring stage development. **a**, PfAP2-HCR-3HA:*loxP* parasites were prepared by U-ExM, stained with N-hydroxysuccinimide (NHS) ester (grayscale), BODIPY FL (white), SYTOX (cyan), and anti-HA antibodies (magenta) and imaged by Airyscan microscopy. The white dashed line marks the nuclear envelope. Scale bars = 5 µm. Images are maximum-intensity projections; numbers indicate the Z-axis thickness of the projection in µm. **b**, Giemsa-stained smears prepared at different time points, from 8 hours post-invasion (h.p.i.) in growth cycle 0 up to 110 h.p.i. Rapamycin was added at 8, 24, and 36 h.p.i. to conditionally truncate *pfap2-hcr* at different stages of the intraerythrocytic developmental cycle (IDC), revealing that PfAP2-HCR is primarily required during the ring stage.

**Extended Data Fig. 4.**
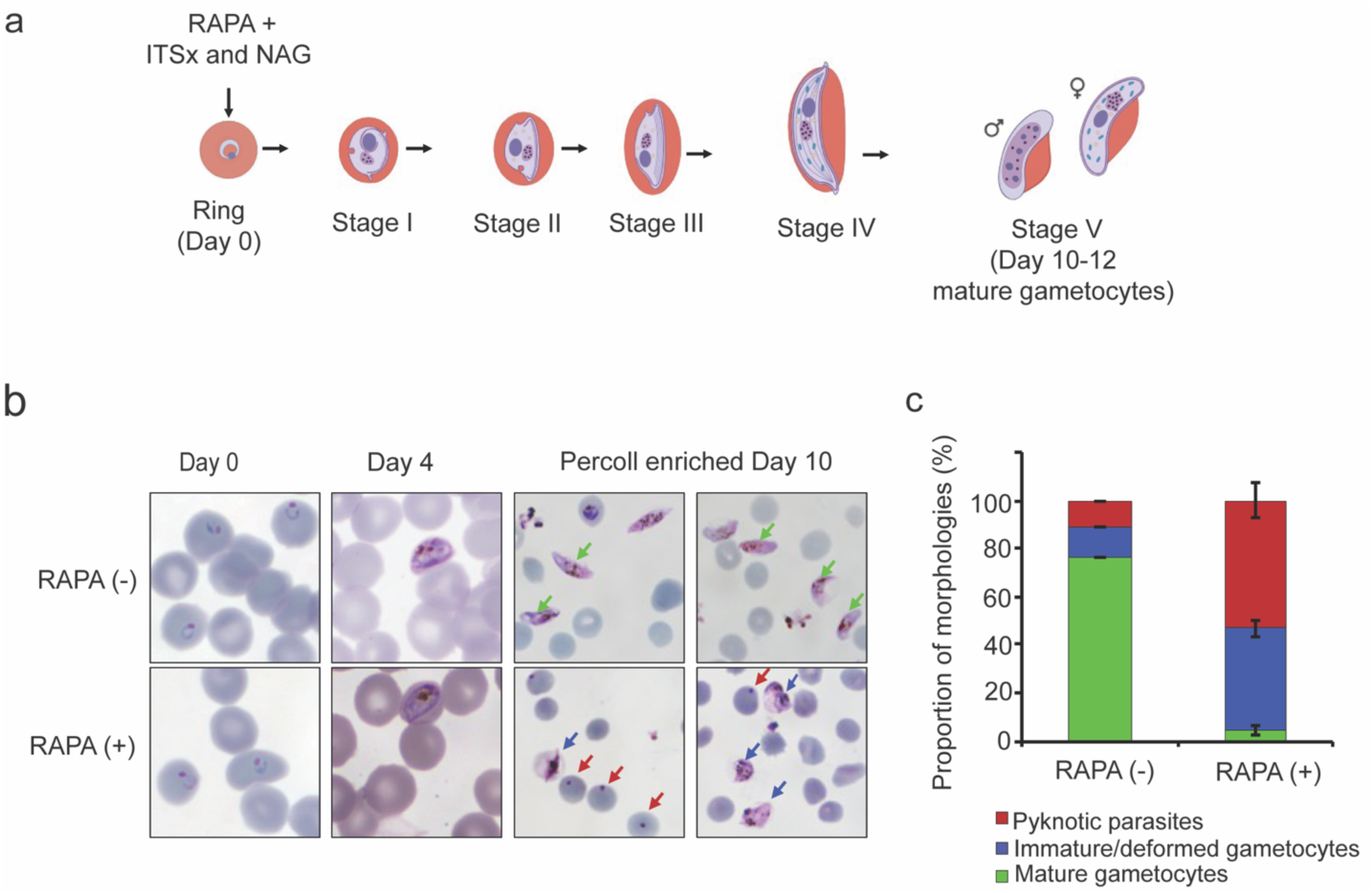
PfAP2-HCR is essential for gametocyte development and maturation. **a**, Schematic representation of the gametocytogenesis induction along with *pfap2-hcr* conditional truncation. **b**, Giemsa-stained microscopic images from day 0, 4 and percoll-enriched day 10 cultures from either DMSO or rapamycin treated parasites. **c**, Quantification of gametocytes (mature/immature/deformed) and pyknotic parasites from DMSO or rapamycin treated day 10 percoll-enriched parasites. Mature, deformed gametocytes and pyknotic parasites are highlighted in green, maroon and blue arrows, respectively. Error bars are representation of standard deviation from two independent experiments.

**Extended Data Fig. 5.**
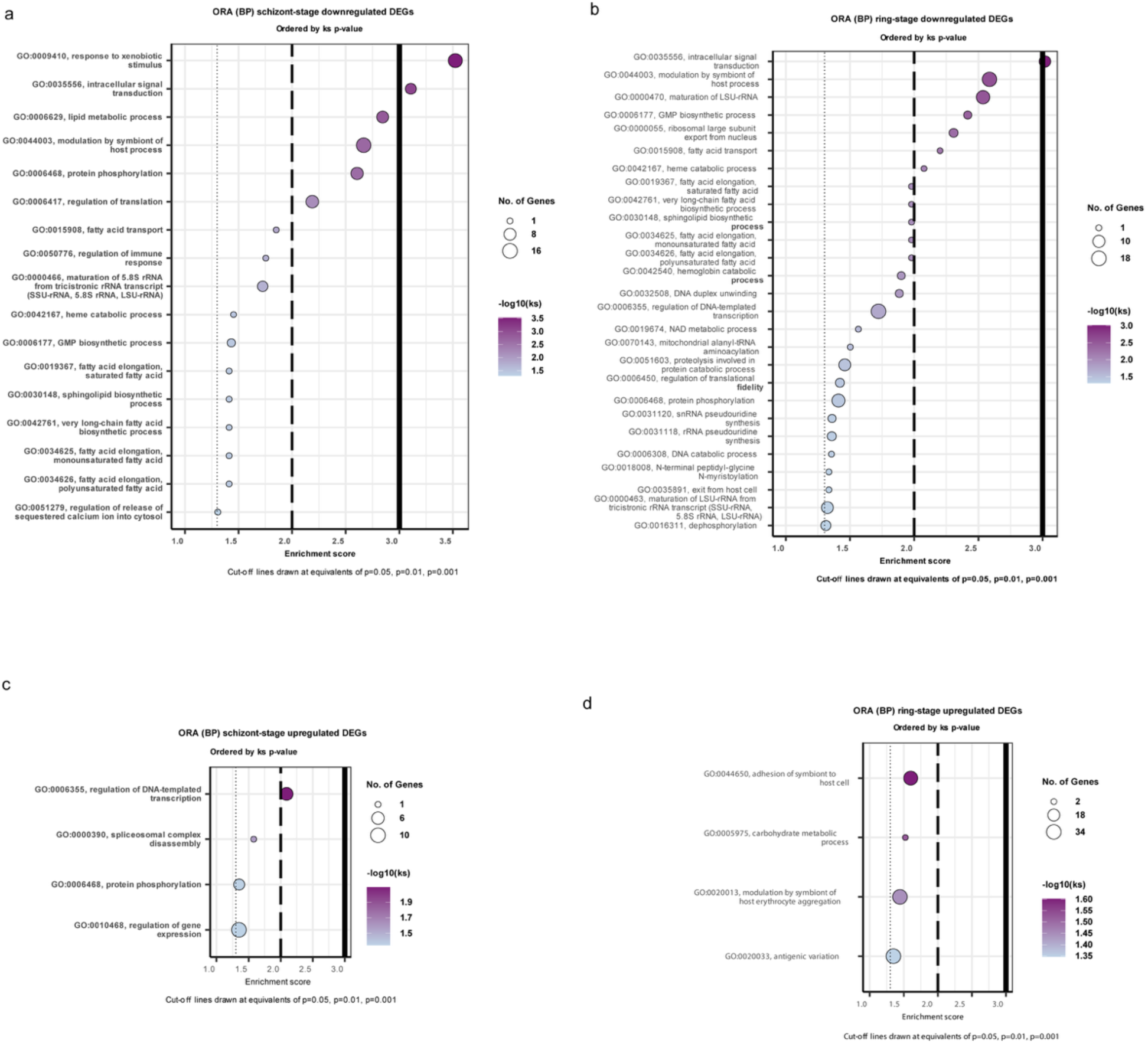
Functional disruption of PfAP2-HCR perturbs multiple biological processes. **a–d**, Gene Ontology (GO) enrichment analysis of biological process terms associated with significantly downregulated (**a, b**) and upregulated (**c, d**) genes in Δpfap2-hcr parasites during the schizont stage (**a, c**) and ring stage (**b, d**).

**Extended Data Fig. 6.**
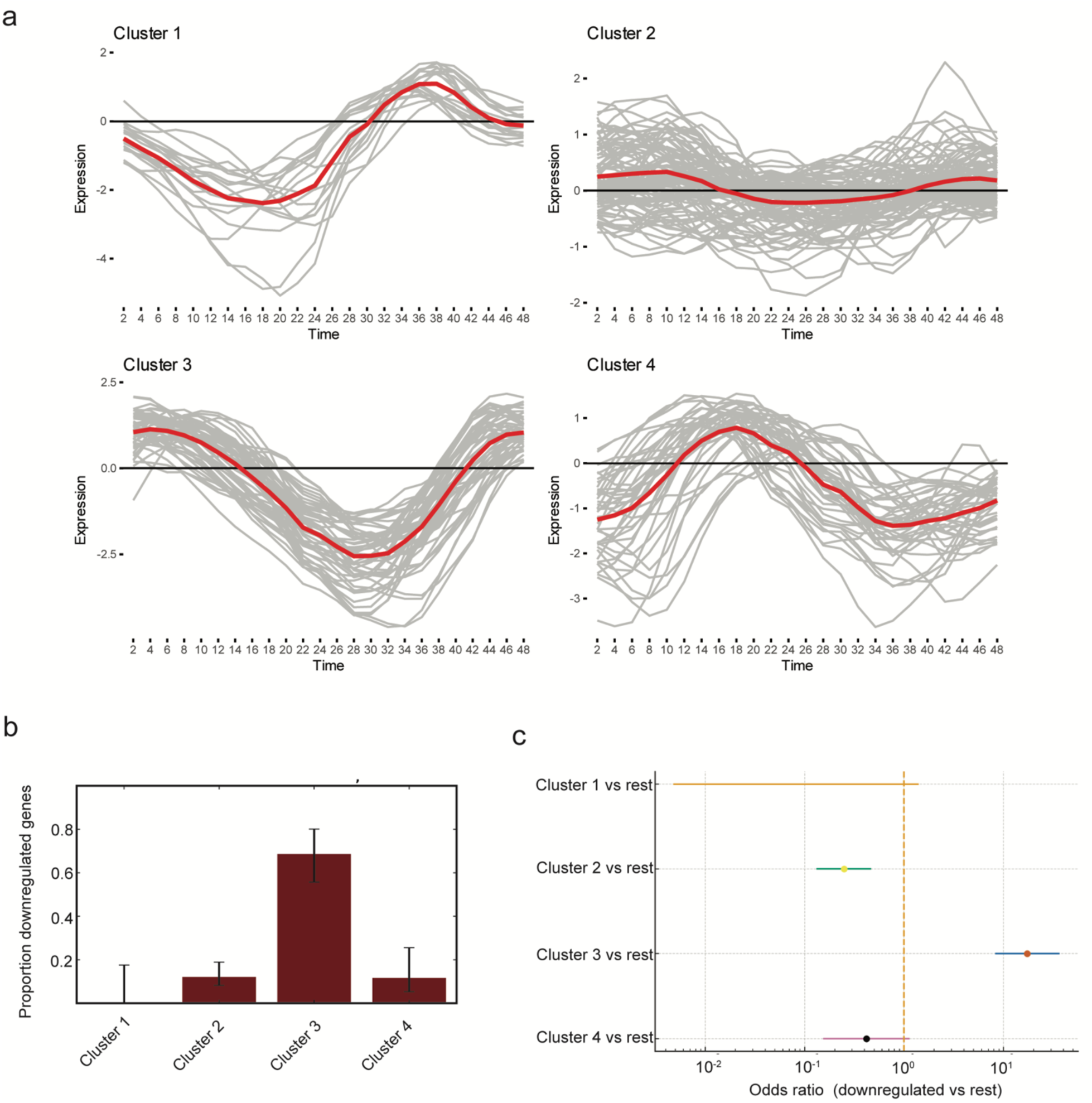
PfAP2-HCR preferentially regulates late schizont and early ring stage exportome genes. **a,** Line graphs showing four expression clusters of exportome based on their expression during IDC. Time in h.p.i. **b**, Fraction of downregulated genes per exportome cluster in Δ*pfap2-hcr* parasites. Error bars indicate 95% Wilson confidence interval. Downregulation was most pronounced in Cluster 3 (Fisher’s exact test, odds ratio = 17.34, *p* = 2.85 × 10⁻¹⁶). **c**, Forest plot showing odds ratios (95% confidence intervals, log scale) for the proportion of downregulated genes in Cluster 3 compared with each other exportome cluster following *pfap2-hcr* truncation. Each point represents the odds ratio from a two-sided Fisher’s exact test; the dashed vertical line indicates no enrichment (odds ratio = 1). Downregulation was significantly enriched in Cluster 3 relative to all other clusters after Holm correction^83^ (*p* < 0.01).

**Extended Data Fig. 7.**
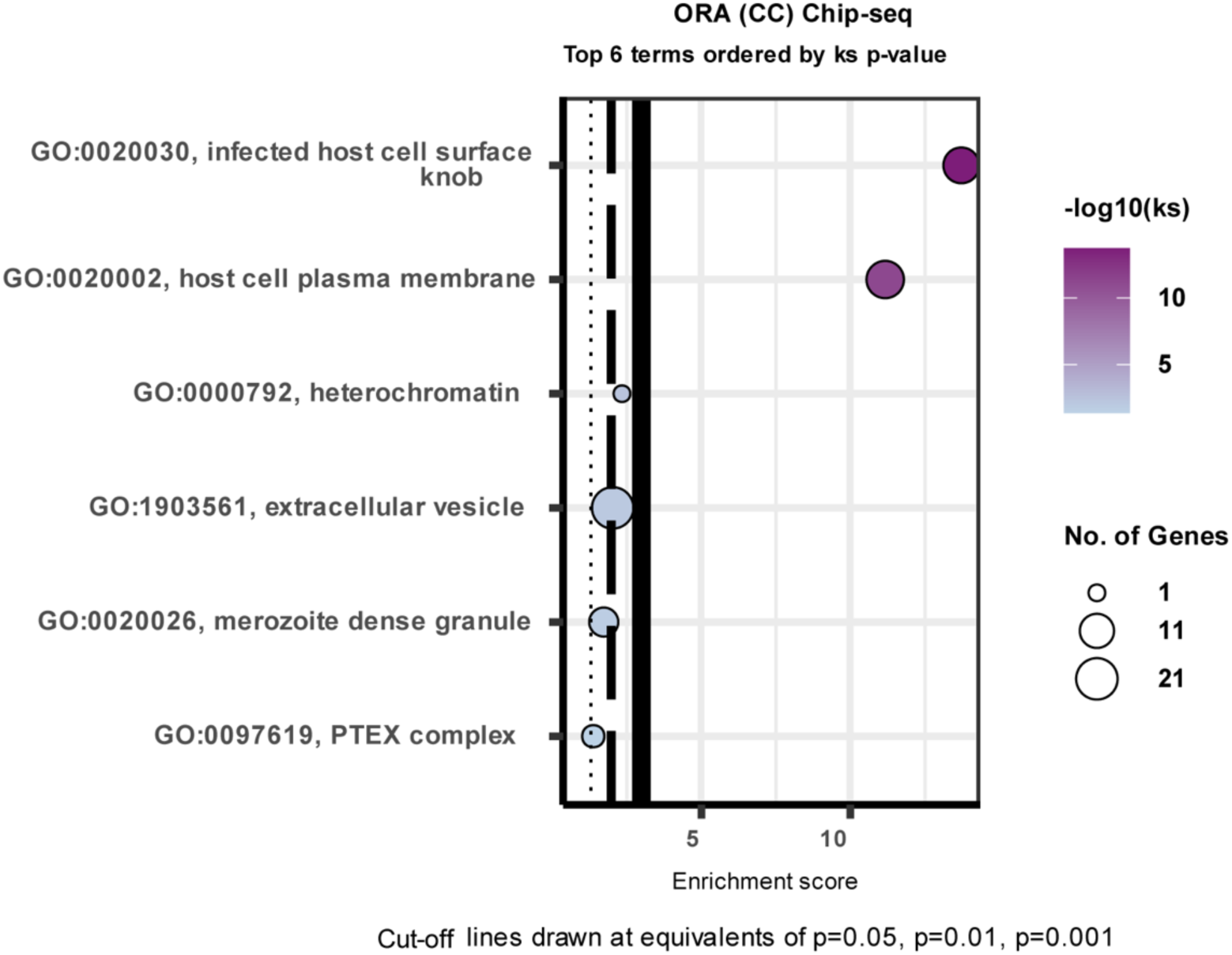
PfAP2-HCR binds to putative promoters of host cell remodeling factors. Gene Ontology (GO) enrichment analysis of cellular component terms associated with genes whose putative promoter regions are bound by PfAP2-HCR, as determined by ChIP–seq. Enriched terms highlight PfAP2-HCR occupancy at loci encoding proteins involved in host cell remodeling, including components of the infected erythrocyte surface, Maurer’s clefts, and the parasitophorous vacuole membrane.

**Extended Data Fig. 8.**
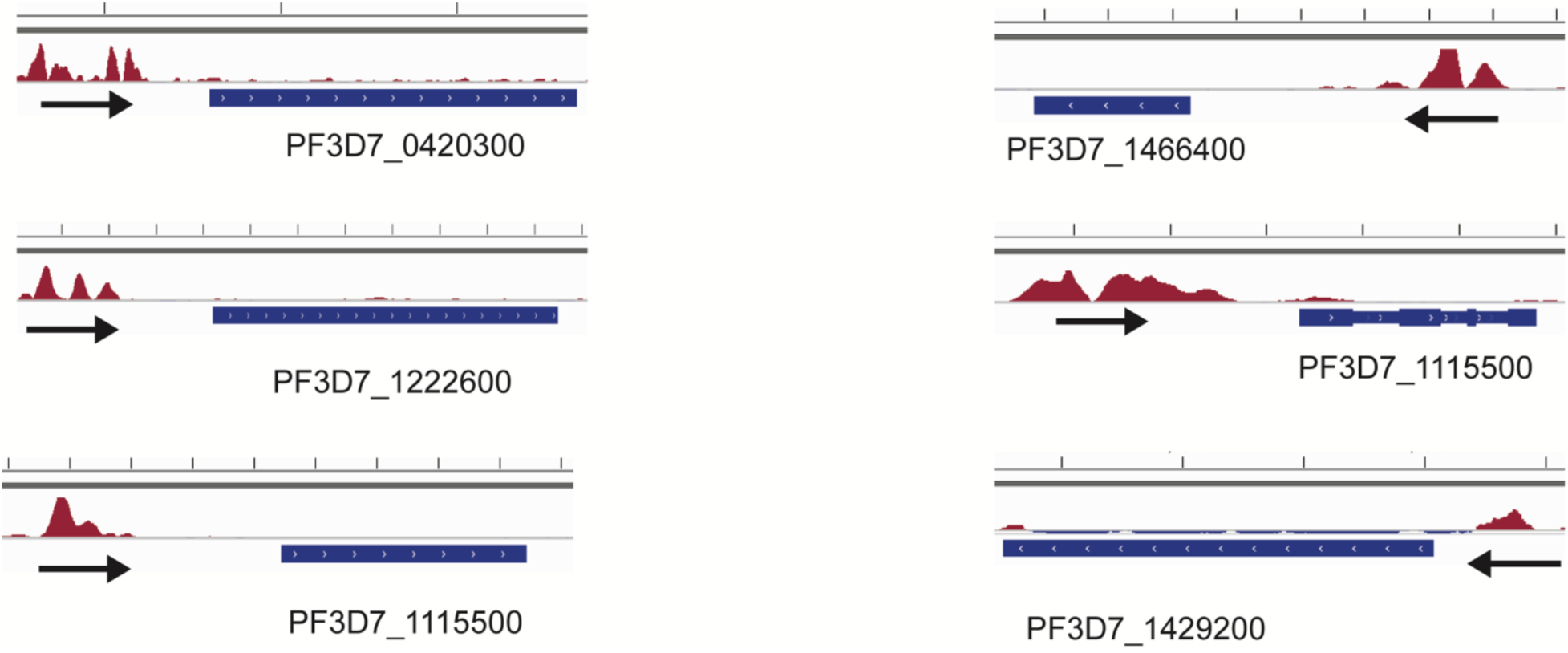
PfAP2-HCR occupancy at the putative promoters of *apiap2* genes. Coverage plots illustrating input-subtracted PfAP2-HCR ChIP–seq enrichment at the putative promoter region of selected *apiap2* genes. Arrows indicate the direction of transcription.

**Extended Data Fig. 9.**
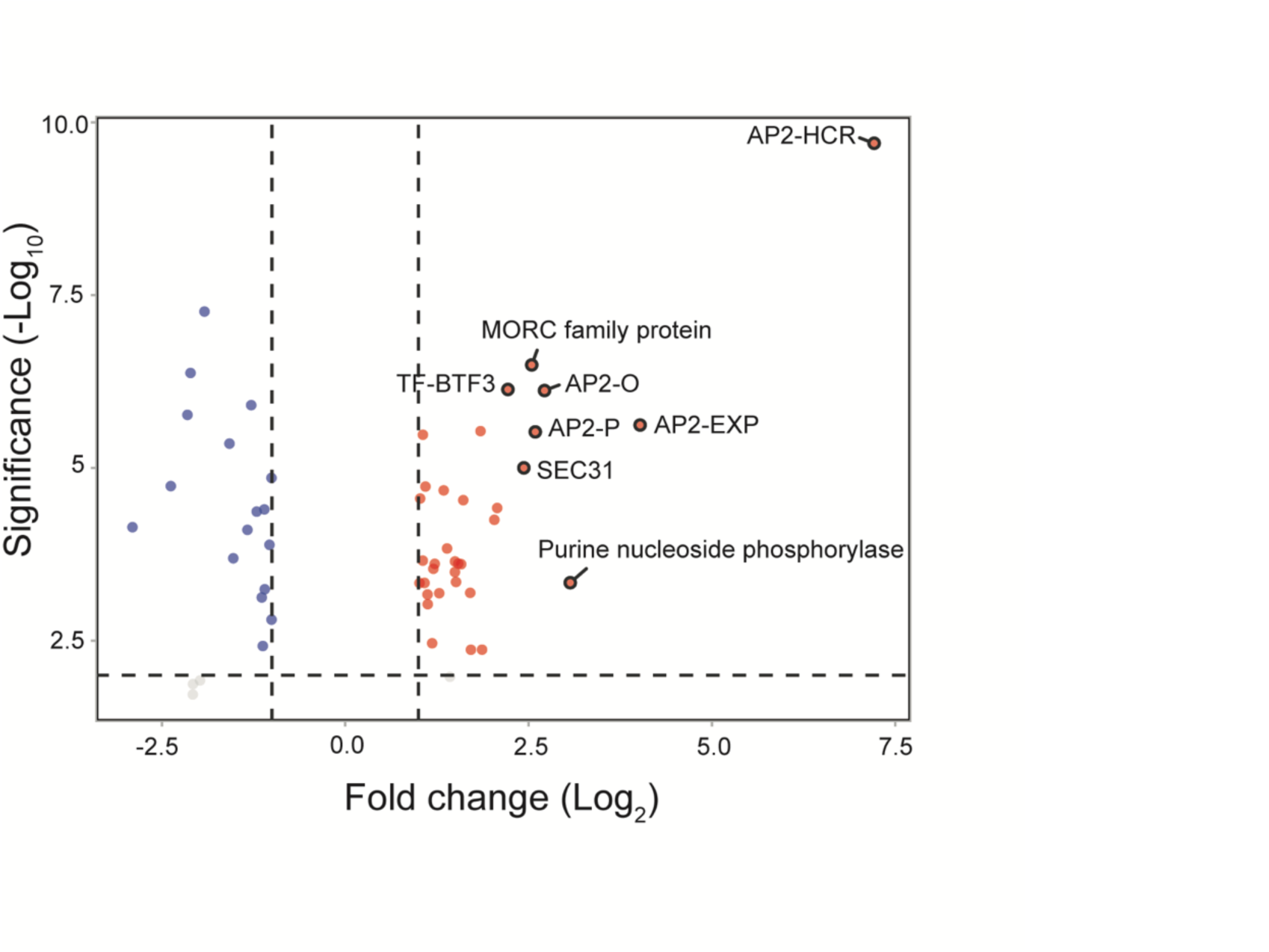
PfAP2-HCR associates with other regulatory proteins. Label-free quantitative proteomic analysis of *P. falciparum* proteins enriched in PfAP2-HCR immunoprecipitates collected at approximately 44 h.p.i. The eight most enriched proteins, ranked by fold-enrichment, are highlighted.

**Extended Data Fig. 10.**
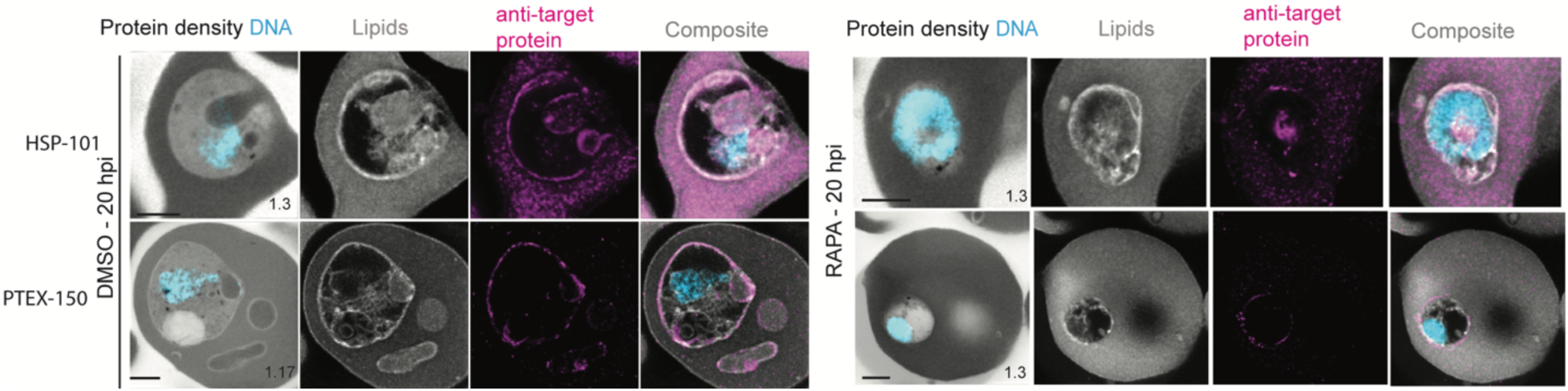
Localization of PTEX components HSP101 and PTEX150 following conditional truncation of *pfap2-hcr*. *Δpfap2-hcr* Parasites were prepared by U-ExM and stained with N-hydroxysuccinimide (NHS) ester (protein density, grayscale), BODIPY FL (lipids, white), SYTOX (DNA, cyan), and antibodies against HSP101 (top) or PTEX150 (bottom) (magenta), and imaged by Airyscan microscopy. In DMSO-treated parasites, HSP101 and PTEX150 display peripheral localization consistent with the parasitophorous vacuole membrane. Images are maximum-intensity projections; numbers indicate the Z-axis thickness of the projection in µm. Scale bars, 5 µm.

## Notes

### Competing Interest Statement

The authors have declared no competing interest.

### Summary of Updates

This revised version of the manuscript has been revised to update the following New data from the gametocytogenesis experiments have been included to demonstrate the essential role of PfAP2-HCR during gametocyte development. The corresponding results are presented in Figure 1h-j and Extended Data Fig. 4a-c.

